# Infant diet promotes *Bifidobacterium* community cooperation within a single ecosystem

**DOI:** 10.1101/711234

**Authors:** Melissa AE Lawson, Ian J O’Neill, Magdalena Kujawska, Anisha Wijeyesekera, Zak Flegg, Lisa Chalklen, Lindsay J Hall

**Author notes:** These authors contributed equally (Melissa AE Lawson, Ian J O’Neill).

## Abstract

Diet-microbe interactions play an important role in modulating the early life microbiota, with *Bifidobacterium* strains and species dominating the gut microbiota of breast-fed infants. Here, we sought to explore how infant diet drives distinct bifidobacterial community composition and dynamics within individual infant ecosystems. Genomic characterisation of 19 strains isolated from breast-fed infants revealed a diverse genomic architecture enriched in carbohydrate metabolism genes, which was distinct to each strain, but collectively formed a pangenome across infants. Presence of gene clusters implicated in digestion of human milk oligosaccharides (HMOs) varied between species, with growth studies indicating within infant differences in the ability to utilise 2’FL and LNnT HMOs between strains. We also performed cross-feeding experiments using metabolic products from growth on 2’FL or LNnT for non-HMO degrading isolates, these compounds were identified to include fucose, galactose, acetate and N-acetylglucosamine. These data highlight the cooperative nature of individual bifidobacterial ‘founder’ strains within an infant ecosystem, and how sharing resources maximises nutrient consumption from the diet. We propose that this social behaviour contributes to the diversity and dominance of *Bifidobacterium* in early life and suggests avenues for development of new diet and microbiota based therapies to promote infant health.

## Introduction

The early life developmental window represents a critical time for microbe-host interactions as this is when foundations for future health and wellbeing are established. Colonisation of pioneer microbes shortly after birth represents a key first step in this mutualistic relationship; shaping the developing microbial community, and in turn impacting numerous host physiological processes (1,2). Although the microbiota of adults is complex in nature, the gastrointestinal (GI) tract of full term healthy infants is relatively simplistic, with upwards of 80% of the total microbiota being comprised of the genus *Bifidobacterium* (1). Loss of *Bifidobacterium*, or indeed gain of other species during this critical window of opportunity, may significantly alter the entire community with negative consequences for host health (3).

In infants, *Bifidobacterium* can be considered a foundation microbiota member which, due to their abundance, strongly influence the environment, the structure of burgeoning microbial communities and host development (4,5). Infant diet is suggested to be one of the key factors that shapes the early life microbiota, and recently the WHO (6) and the Scientific Advisory Committee on Nutrition (UK) (7) released new guidelines regarding the optimal time to start breast feeding, and highlighted the health benefits associated with solely breast-feeding infants. In breast-fed infants, the early life microbial community is dominated by a single bacterial genus *Bifidobacterium;* that is, until the diet changes to include either infant formula and/or simple solid foods. It is known that breast-fed and formula-fed infants differ in microbial composition (8), including significant differences in bifidobacterial populations, which has also been linked to differential health outcomes, both in the short- and longer-term i.e. induction of allergies, asthma and obesity in formula fed infants (8,9).

Establishment of this bifidobacterial dominant community in response to diet (specifically breast milk) suggests a cooperative relationship between different strains and species, rather than competition, which is often described for other bacterial species (10). Indeed, it appears that closely related *Bifidobacterium* strains coexist in a single infant GI tract, rather than one strain dominating and competitively excluding other strains. However, a cooperative balance between bifidobacterial strains in the early life microbiota may further enhance their dominance in breast fed infants by enabling a genus-specific exploitative competition i.e. depleting the GI tract of breast milk-derived nutrients, thereby preventing colonisation of other microbes, including pathobionts. To investigate these key community dynamic questions, we have probed the genomic and phenotypic similarities between bifidobacteria strains that coexist in the same individual, including their responses to specific early life diet components, namely human milk oligosaccharides (HMOs). By examining microbial interactions on a strain-level we provide important insights into how multiple species of *Bifidobacterium* co-exist within a single infant in early life, which may have implications for design of diet- and microbial-based early life therapies.

## Results

### *Bifidobacterium* dominates the infant microbiota

To investigate bifidobacterial community interactions in early life, the faecal microbial community profiles from three full term, healthy infants (referred to as infant V1, V2 and V3) was subjected to metataxonomic profiling using 16S rRNA gene sequencing (Fig 1A and S1A). At the time of sample collection, all infants were similar in age (mean = 145 ± 38 d), born vaginally and exclusively breast-fed (with the exception of two isolations coming from infant V3 at an earlier time point, Fig S1B and Table S1). In agreement with previous studies, we observed a high prevalence of *Bifidobacterium* in each infant faecal sample (mean = 82.53±12.36%). Further analysis indicated a dynamic bifidobacterial community comprised of different strains and species present in each infant, thus we isolated multiple *Bifidobacterium* strains to explore bifidobacterial community dynamics. A total of 19 strains were isolated and their genetic relatedness was inferred based on core genome phylogeny (Fig 1B), and Average Nucleotide Identity (ANI) values, and compared to 83 publicly available bifidobacterial genomes (Fig S2). The generated phylogenetic tree indicates the clustering of all analysed strains into three main phylogenetic groups denoted as the *Bifidobacterium longum* (encompassing the members of the *longum* and the *infantis* subspecies), *Bifidobacterium breve* and *Bifidobacterium pseudocatenulatum* groups. Strains isolated from infant V1 were classified as either *B. longum* subspecies *longum* (hereafter referred to as *B. longum*) or *B. pseudocatenulatum*; V2 strains classified as *B. breve* or *B. longum* subspecies *infantis* (hereafter referred to as *B. infantis*), and V3 strains were classified as either *B. pseudocatenulatum, B. infantis*, or *B. longum* (Fig 1B). Genome sizes ranged from 2.25Mb (*B. pseudocatenulatum* LH13) to 2.75Mb (*B. longum* subsp. *infantis* LH23), with an average of 2.38Mb, consistent with previously published data (11). The G+C content ranged from 56.50% for *B. pseudocatenulatum* LH9 to 60.04% for *B. longum* subsp. *longum* LH277, while the number of predicted ORFs was lowest in *B. pseudocatenulatum* LH11 (1,888), and highest in *B. longum* subsp. *infantis* LH23 (2,521) (Table S1).

**Figure 1:**
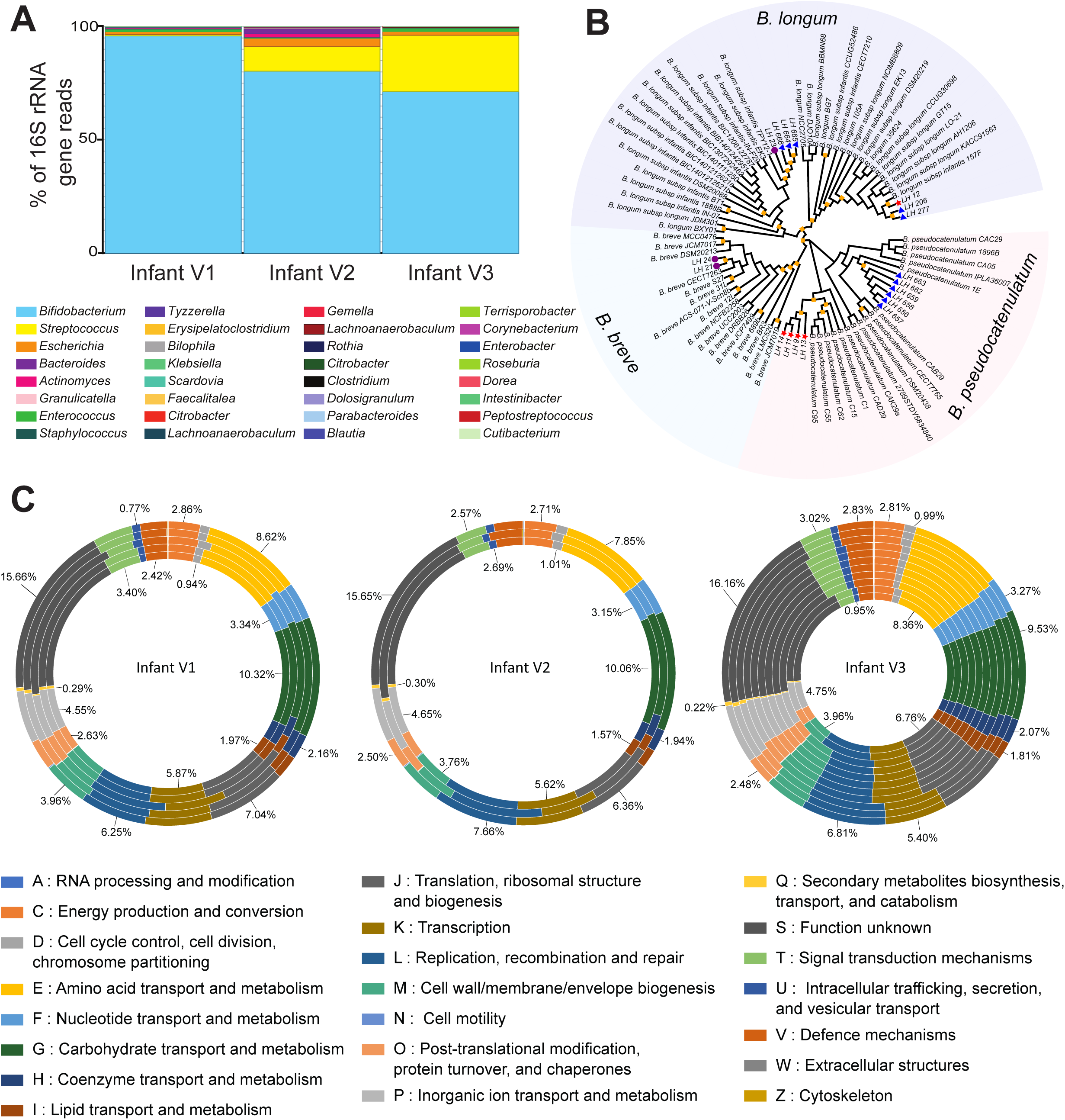
Diverse *Bifidobacterium* strains dominate the early life microbiota. (A) Faecal bacterial community profiles of three healthy, full-term infants as determined by 16S rRNA gene sequencing. Paired-end reads were generated using the MiSeq Illumina platform, all data sets were normalised and relative abundance of each bacterial taxa is represented in percentages of number of total reads for the top 10 most prevalent genus in each infant. Bar colours represent different genus taxa, and bar lengths signify the relative abundance of each taxon. 16S rRNA bacterial profiles are named according to the sample used for bifidobacteria isolation. V1 at 102 days of age, V2 at 174 days of age, and V3 at 159 days of age. The number of reads obtained by 16S rRNA gene sequencing data for each sample can be found in Table S2. (B) Core-genome phylogeny of 83 *Bifidobacterium* isolates, 19 of which are novel strains identified in this study and denoted by arrows and the annotation of LHXX. Isolates from infant V1 are denoted with a red star, V2 with a purple circle and V3 with a blue triangle. Bootstrap values >70 are shown with a yellow square on each node. (C) ORFs from each genome was submitted to eggNOG-mapper (http://eggnogdb.embl.de/#/app/emapper) for functional classification. The proportion of ORFs for each classification was calculated and is presented as a percentage of total ORFs in each genome. The values indicated for each orthologous group represents the average percentage of ORFs in that group for all genomes in that infant.

### Analysis of bifidobacterial genomes reveals distinct and diverse functional profiles within and between infant isolates

Previous studies have highlighted that diet may act as an evolutionary driver in *Bifidobacterium*, as functionally, 13.7% of genes in the bifidobacterial pan genome are involved in carbohydrate metabolism, thereby reflecting their ability to grow within the glycan rich environment of the colon (12,13). We functionally characterised genomes and determined that, apart from genes of unknown function, genes classified as carbohydrate transport and metabolism were the most abundant in all genomes, indicating the saccharolytic lifestyle of *Bifidobacterium* spp. (Fig 1C and Table S3). Strains from infant V1 had the highest percentage of carbohydrate metabolism and transport genes of all infants (10.32%), followed by strains from infant V2 (10.06%), and then infant V3 (9.53%). Based on functional genome analysis there appears to be interspecies differences in genes associated with carbohydrate transport and metabolism, for example, 10.67± 0.11% of the total genome of *B. pseudocatenulatum* strains from infant V1 were dedicated to this functional group, whereas V3 *B. pseudocatenulatum* strains had 1% less (9.60±0.18%) of their genome annotated for carbohydrate use. These data suggest the same species in different ecological niches (i.e. infant) may have altered metabolic function.

Due to the high proportion of *B. pseudocatenulatum* strains isolated from infant V1 and V3, and apparent differences in functional annotation, we further investigated differences in the core and accessory genes (Fig. S3A, Tables S6A & S6B). *B. pseudocatenulatum* strains from the same infant shared a large core genome, but there were minor differences in accessory genes. Strains from infant V1 had more unique genes compared to strains from infant V3 (Fig S3). Further characterisation of these unique genes indicated that infant V1 strains LH13 and LH14 possess unique putative carbohydrate utilisation gene clusters (LH_13_00067-LH_13_00071 and LH_14_01835-LH_14_01839) that are not present in other strains and may contribute to altered carbohydrate digestion. The LH14 genome also contains genes encoding fimbriae and sortase genes (LH_14_01074-LH_14_01081), suggesting presence of sortase dependent fimbriae. We have also identified a partial prophage gene cluster in LH13 (LH_13_01645-LH_13_01657) that encodes phage tail and head proteins and a tyrosine recombinase, however, due to the size of the cluster it is likely to be an unsuccessful phage insertion event. Interestingly, two large partial prophage clusters were also identified in genomes of LH656 and LH658 from infant V3 (Table S7A & S7B).

*Bifidobacterium* possess a large repertoire of glycoside hydrolases (GH) that facilitate digestion in the glycan rich gut environment, therefore, we determined the number of GH present in each infant faecal ecosystem (Fig 2 and S3B). Genes from a total of 39 different GH families were represented in at least one genome with *B. pseudocatenulatum* strains have the highest number of GH genes (mean = 62 per genome), followed by *B. longum* (mean = 48 per genome), *B. breve* (mean = 46 per genome), and *B. infantis* (mean = 42 per genome). In all infants, the most represented family were members of GH13 representing enzymes involved in the hydrolysis of alpha-glucosidic linkages found in plant di-, oligo- and polysaccharides. The second most abundant GH family in infants V1 and V3 (but not V2) was GH43 which contains beta-xylosidases involved in the digestion of the plant polysaccharide xylan. Other highly represented families were GH3, containing beta-glucosidases that hydrolyse a wide range of glycans present in plant cell walls, GH2 and GH42, containing beta-galactosidases that are active on galactooligosaccharides and galactans found in plant cell walls, and also active on lactose that is present in human breast milk.

**Figure 2:**
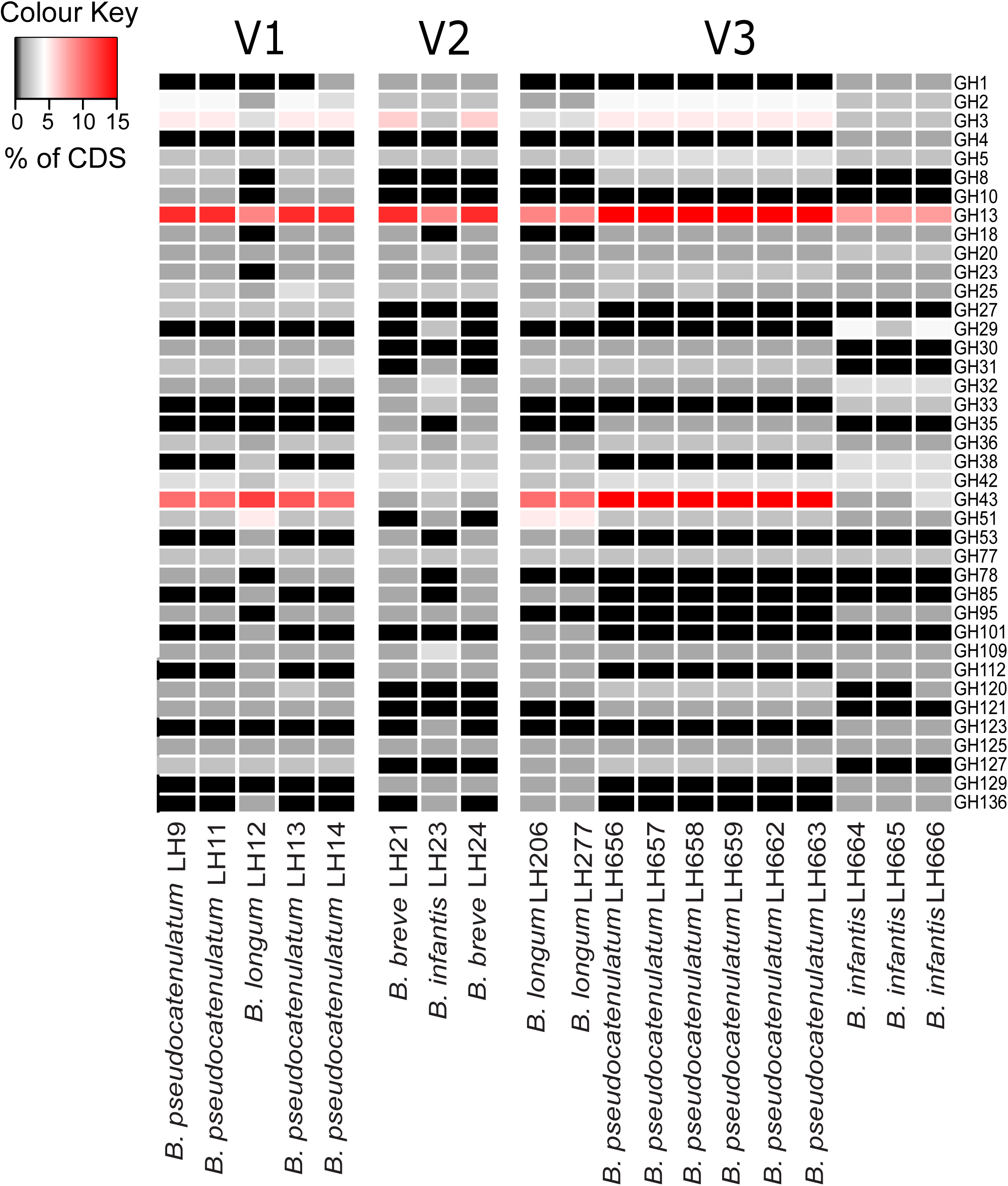
Functional classification of *Bifidobacterium* genomes. (A) Presence of genes encoding glycosyl hydrolases was determined using the dbCAN server (http://csbl.bmb.uga.edu/dbCAN/) which annotates genes based on HMMs of GH generated from data in the CAZy database (http://www.cazy.org). The heatmap shows the number of ORFs annotated as GH for each GH family (y-axis) for each genome (x-axis) (Enumeration of GH ORFs in subfamilies of GH5, GH13 and GH43 is shown in Supplementary figure 3B).

Mucin present in the infant gut contains sialic acid residues that can be utilised by many members of the microbiota including *Bifidobacterium* (14). *B. infantis* and *B. brev*e strains from infants V2 and V3 contains exo-sialidases from GH33 potentially allowing these strains to utilise host sialic acids (Fig. 2). Interestingly *B. pseudocatenulatum* and *B. longum* from infant V3 do not contain these genes, however, crossfeeding of sialic acids between *Bifidobacterium* species has previously been reported (15).

Ultimately, the presence of multiple strains within the *Bifidobacterium* community results in a larger repertoire of GHs, potentially indicating an ability to digest a wider range of oligosaccharides within the infant gut.

### Infant bifidobacterial isolates have enhanced survival against exposure to intestinal environmental stressors

To colonise and maintain a successful bifidobacterial ecosystem, strains must first be able to traverse the length of the GI tract, to their preferred niche i.e. the colon. When exposed to acid shock at pH2 for four hours (representing stomach pH and transit time), all strains (except *B. breve* LH24) were able to re-establish growth in MRS media (Fig 3A). As expected all strains exhibited poor growth in aerobic, compared to anaerobic conditions, although there was modest growth in some strains, which might suggest an ability for vertical and horizontal transmission (Fig 3B). Growth in 0.3% bovine bile salt resulted in statistically significantly less growth of all strains, except for *B. infantis* LH23 from infant V2 (Fig 3C). All strains contained homologs to known bile salt hydrolase genes (Table S3), and therefore we tested the ability of each strain to hydrolyse specific individual bile salts including taurocholic acid, taurodeoxycholic acid and sodium glycodeoxycholate bile acids (Table S4). We found differences between strains from the same individual, such as *B. pseudocatenulatum* LH11 was capable of hydrolysing taurocholic acid, taurodeoxycholic acid and sodium glycodeoxycholate bile acids, while *B. longum* isolates (LH206 and LH277) were unable to hydrolyse any of the bile salt tested, and only the *B. longum* strain from infant V1 (LH12) could precipitate taurodeoxycholic bile acid. Within a single infant bifidobacterial community, it appears there are those strains that have enhanced intestinal stressor survival, compared to highly related intra-infant strains, indicating adaptation that may enhance colonisation potential of these strains, thus impacting community succession.

**Figure 3.**
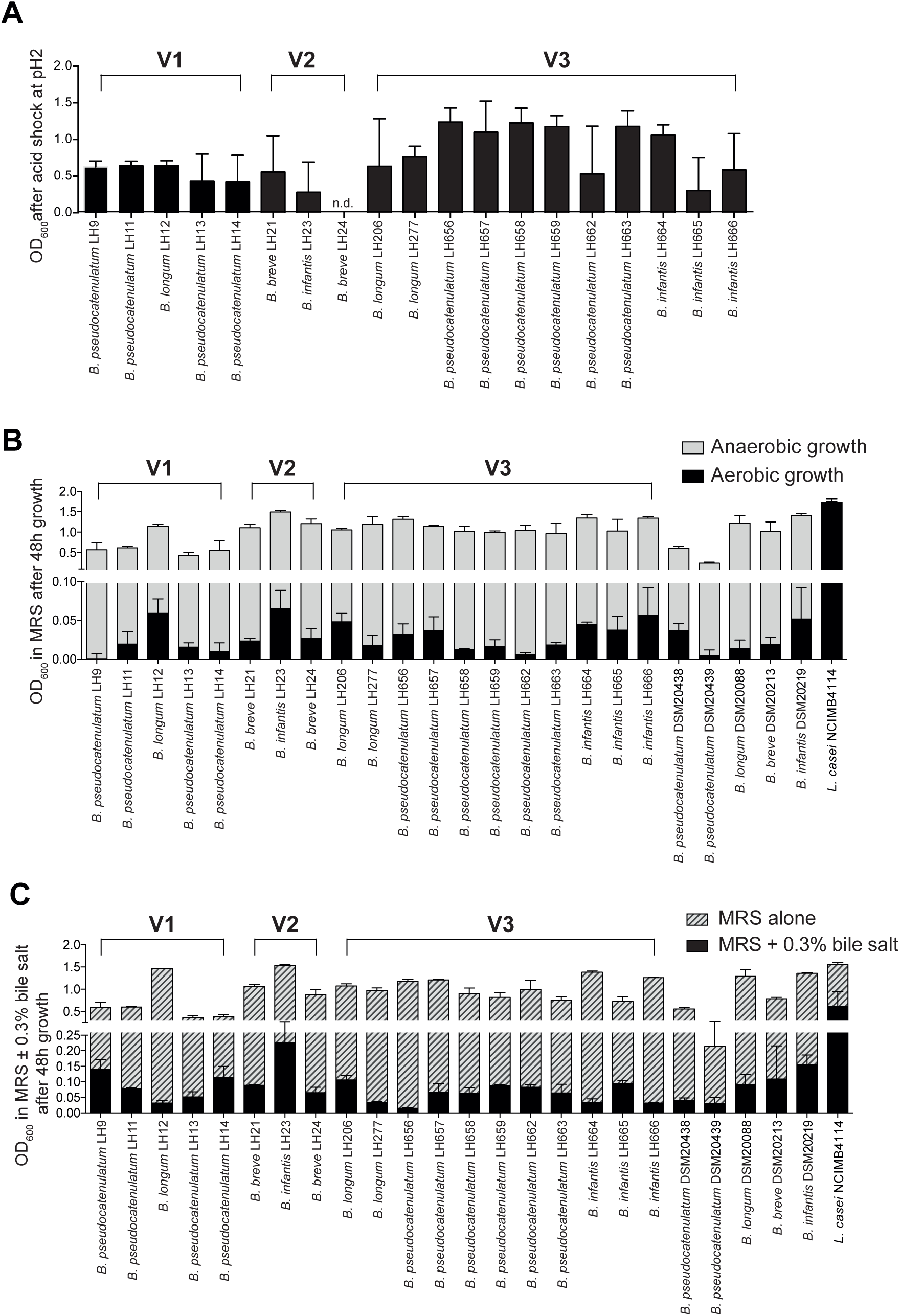
Bifidobacterial survival after exposure to intestinal environmental stressors. (A) Bacterial growth in MRS was measured for each strain after 30h growth in an aerobic environment and compared to parallel samples grown anaerobically. Measurements shown are representative of three independent experimental repeats. (B) Ability to withstand acid shock for 4 h at a pH2. Following acid shock, strains were incubated for 48 h in an anaerobic cabinet at 37°C. Growth (optical density) was then measured. Data shown is mean ± standard deviation for 3 independent experiments (C) Bacterial survival in MRS supplemented with 0.3% bile salts was measured after 48h growth and compared to parallel cultures grown in MRS anaerobically at 37°C. Data shown from five independent experiments; n.d. denotes measurement not detectable. Blank/media only values are subtracted from all points in each experiment, all negative values are represented as zero. Isolates from Baby V1 are represented as black bars, V2 are grey bars, V3 are dotted bars, and the tested type and control strains are shown as white bars. Error bars represent SEM.

### The presence of HMO gene clusters is species and strain specific

With respect to an infant-specific diet, *Bifidobacterium* genomes encode a large repertoire of glycosyl hydrolases (GH) that allow for digestion of human milk oligosaccharides (HMO) (Katayama, 2016), which are a key component of maternal breast milk. To date, there have been over 200 different HMOs identified in maternal milk, which are unconjugated glycans comprised of a lactose core with varying chain length of 3 to 15 carbohydrates containing either glucose, galactose, fucose, N-acetylglucosamine (GlcNAc), and N-acetylneuraminic acid (NeuAc) or sialic acid (16,17). In addition, the lactose backbone of HMOs can be either fucosylated or sialylated to form trisaccharide HMO structures, namely 2’ or 3’-fucosyllactose (FL) and 3’ or 6’-sialyllactose (SL), respectively. Gene clusters involved in the digestion of HMOs have also been described for *B. infantis, B. breve, B. longum* and *B. pseduocatenulatum*, however, the genetic organisation and size of these HMO clusters varies from species to species (18–22). Thus, we analysed the genomes in this study for the presence of the clusters previously described for *B. infantis, B. breve, B. longum* and *B. pseduocatenulatum.*

Comparison with the large, 45kb, *B. infantis* ATCC 15697 HMO cluster (BLON_RS12070-BLON_RS12215) identified similar clusters in *B. infantis* LH23, LH664, LH665 and LH666 (Fig 4A), however, the genetic organisation of this cluster varied in our infant isolates. Importantly, homologues to all three GH contained within the cluster are found in similar clusters in our genomes, however, instead of a large continuous cluster, homologues of genes from ATCC 15697 are found in two separate clusters within LH23, which may reflect the presence of transposon sequences flanking each cluster. *B. infantis* LH665 lacks four transport-reported permease genes present in the ATCC 15697 cluster, however, other transport binding proteins and permeases encoded within the LH665 HMO cluster may perform a similar function.

**Figure 4.**
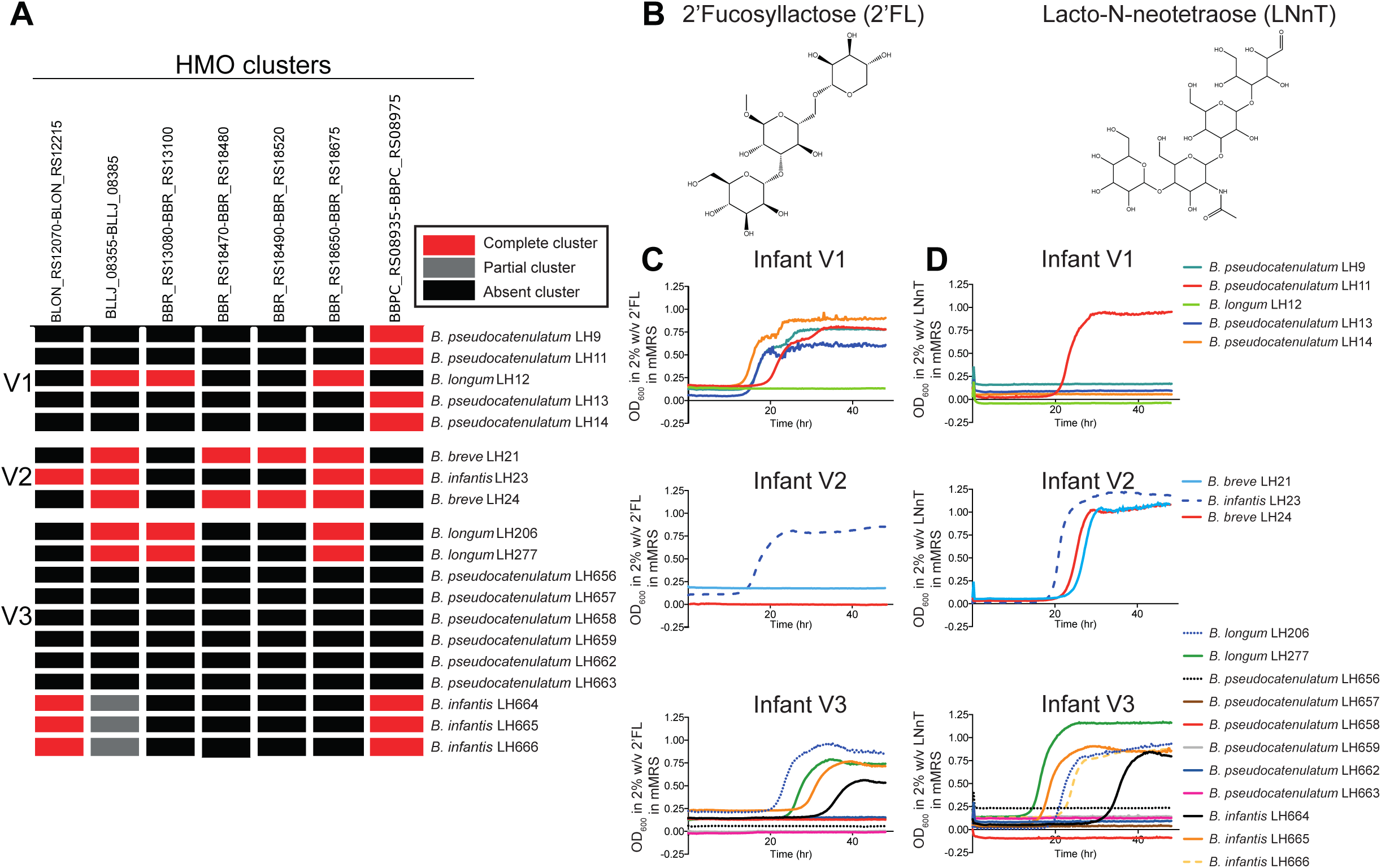
HMOs function as key carbon sources for *Bifidobacterium* growth. (A) HMO heatmap illustrating the presence of known HMO gene clusters in the 19 novel infant isolates (see Methods for details); (B) chemical structure of HMO 2’FL and LNnT generated by ChemDraw; (C-D) Growth kinetics of all 19 strains in mMRS with either HMO 2’FL (C) or LNnT (D) as a sole carbon source; all strains from each infant are illustrated together. Data shown is a representative graph of three independent experimental repeats, containing the mean from duplicate/triplicate well measurements.

Digestion of 2’FL in *B. longum* SC569 is linked to the presence of genes encoding an alpha1-3/4-fucosidase (GH29) and/or alpha1-2 fucosides (GH95), which are present in a gene cluster with carbohydrate utilisation genes (BLNG_01254-BLNG_01264). Interestingly, neither this cluster nor any alpha-fucosidades were identified in any of the *B. longum* strains from any of the infants. Homologues to these GHs were identified in all *B. infantis* (LH23, LH664, LH665 and LH666) strains from infant V2 and V3 (data not shown). A highly similar gene cluster is found in *B. pseudocatenulatum* DSM 20438, but only contains one alpha-fucosidase (GH95) and all *B. pseudocatenulatum* strains in infant V1, LH9, LH11, LH13 and LH14, encode homologs of these clusters, including the key GH95 gene (Fig 4A).

*B. breve* UCC2003 encodes four separate gene clusters; *lnt* cluster (BBR_RS13080-BBR_RS13100); *lac* cluster (BBR_RS18470-BBR_RS18480), the *nah* cluster (BBR_RS18490-BBR_RS18520) and *lnp/glt* cluster (BBR_RS18650-BBR_RS18675) that are important for the digestion of LNT and LNnT (22,23). Homologs to the *lac, nah* and *lnp/lgt* clusters were identified in *B. breve* LH21 and LH24. The *lnt* cluster is upregulated during growth on LNT and LNnT, but this cluster was not present in either LH21 or LH24, however, it was identified in B. *infantis* strains LH206 and LH277, and in *B. longum* strain LH12. Unsurprisingly, the *lnp/glt* cluster was identified in *B. longum* strains LH12, LH206, LH277 as this cluster has also been described for *B. longum* JCM1217. *B. infantis* strains LH664, LH665 and LH666 contained a partial *lnp/glt* cluster with highly similar enzymatic genes, but no transporter genes in close genomic proximity, suggesting that these strains may degrade LNB, but may transport this HMO using other genes in the genome.

Sialidases aid breakdown of sialidated HMOs and *B. bifidum* ATCC 15696 has an extracellular sialidase (SiaBb1) that can digest these HMOs without importing them into the cytosol (24,25). Our analysis identified sialidases in *B. infantis* strains LH23, LH664, LH665 and LH666, however, the required transmembrane domains were absent suggesting that these strains are intracellular sialidase utilisers (Table S8). Thus, it appears that infant-specific strains may maximise utilisation of a wide assortment of HMOs within a community via differential genetic adaptation to a breast-milk diet and early life niche.

### Phenotypic characterisation of HMO usage indicates known and unknown enzymatic gene clusters

Next to lactose, HMOs are the second most abundant carbohydrate in breast milk (5-15 g/L in breast milk) (26), and are a key metabolic source for *Bifidobacterium* (27,28). Although we identified a wide range of putative HMO genomic clusters in our strains, we next sought to correlate these with an ability to metabolise HMOs, in particular 2’FL and LNnT (Fig. 4B-D). We were able to identify strains capable of using either 2’FL or LNnT in each microbial ecosystem. All *B. pseudocatenulatum* strains, but not the *B. longum* LH12 strain, from infant V1 could degrade the HMO 2’FL, most likely due to the presence of a known fucosylated HMO utilisation gene cluster (Fig 4A). Only *B. pseudocatenulatum* LH11 was capable of using the more complex HMO, LNnT, despite lacking known enzymatic clusters, suggesting an alternative or novel mechanism of degradation. Interestingly, only strain *B. infantis* LH21 isolated from infant V2 could degrade 2’FL for growth (neither *B. breve* strains LH23 or LH24 could), and yet all three strains grew well with LNnT as a sole carbon source. None of the *B. pseudocatenulatum* strains from infant V3 could use either of the HMOs tested, possibly due to the absence of appropriate HMO clusters in their genomes, while all the *B. longum* and *B. infantis* strains appeared to metabolise both HMOs for growth; apart from *B. infantis* LH666 which only utilised LNnT for growth. This data is supported by bioinformatic analysis identifying a large HMO degradation gene cluster and the presence of key alpha-fucosidases in all *B. infantis* strains. Collectively these data demonstrate that HMO utilisation is dependent on the type of HMO and the strain tested; supported by the fact that type strains for each species also had differential HMO degradation (Fig. S4).

### A syntrophic network for HMO utilisation exists within *Bifidobacterium* species

It is clear that multiple stains and species exist as a community within a single ecosystem (i.e. infant), and that these bifidobacterial communities encode a diverse array of metabolic genes, which correlates with strain-specific breakdown of breast milk-associated HMOs. Thus, to assess how these bifidobacterial communities interact with each other, we determined if strains unable to degrade HMOs, could utilise the metabolic by-products generated by HMO degraders. To address this experimentally, all strains that had previously been found to use either 2’FL (Fig 4C) or LNnT (Fig 4D) were grown in their respective HMOs, and conditioned media was filtered and used as a carbon source for all other non-HMO users within the same infant community (Fig 5A). We found that 2’FL derived-substrates from all *B. pseudocatenulatum* strains in infant V1 supported growth of *B. longum* LH12 (Fig 5B). This demonstrates a syntrophic/cross-feeding network, and cooperation between different species within the same ecological niche. Conversely, V2-associated *B. breve* strain breakdown products did not support *B. infantis* LH23 growth, indicating that these strains were not able to cross-fed when HMO 2’FL is the initial carbon source. Both *B. longum* and two *B. infantis* isolates from infant V3 (LH206, LH277, LH664 and LH665 respectively) grew on 2’FL, however only the conditioned media from isolate *B. longum* LH206 could support growth of other isolates within the same infant. Interestingly, bioinformatic analysis did not identify any alpha-fucosylase genes in LH206. Moreover, LH206 conditioned media enhanced growth of all tested isolates (both *B. infantis* and *B. pseudocatenulatum* species), which suggests the metabolism of 2’FL by LH206 may generate a wide variety of components for neighbouring isolates to use for growth within this ecosystem (i.e. infant V3).

**Figure 5.**
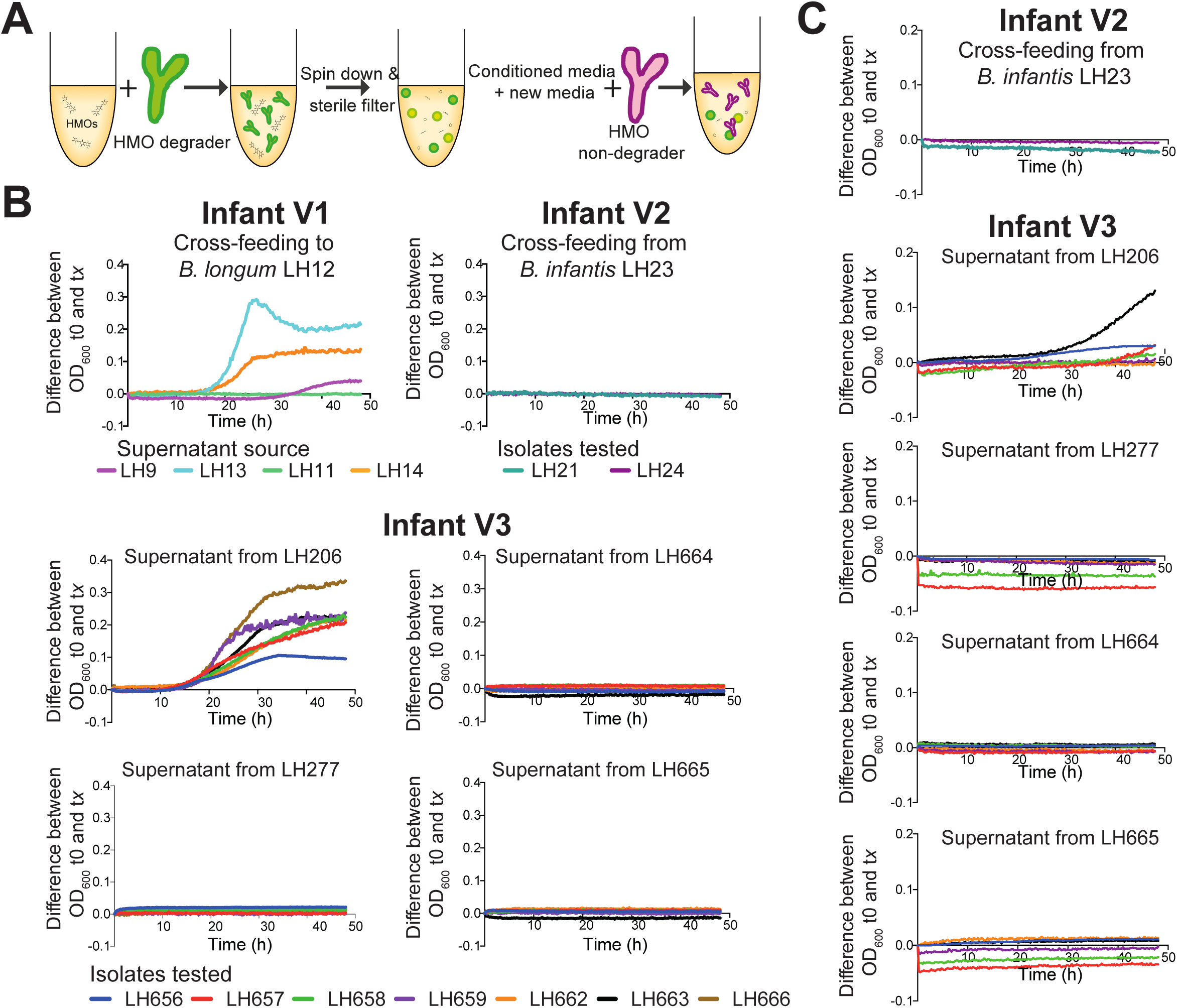
Cross-feeding networks exist within *Bifidobacterium* communities. (A) Schematic illustrating experimental set-up and preparation of conditioned media for cross-feeding experiments. (B) Using conditioned media from strains capable of using either HMO 2’FL (B) or LNnT (C) was combined with fresh mMRS (1:1) and used as growth media for all HMO non-degrader isolates within the same ecological niche. Data shown is a representative graph of three independent experimental repeats, containing the mean from duplicate/triplicate well measurements. (D)^1^H NMR spectrum to identify metabolites involved in cross-feeding between 2’FL-degrader *B. longum* LH206 and non-degrader *B. pseudocatenulatum* LH659 and semi-quantification of identified metabolites. (E) ^1^H NMR spectrum to identify metabolites involved in cross-feeding between LNnT-degrader *B. longum* LH206 and non-degrader *B. pseudocatenulatum* LH657 and semi-quantification of identified metabolites. n.d indicates metabolites were not detected.

Overall fewer isolates could grow using the more complex HMO LNnT, and as a result no strains from infant V1 were tested. In contrast to the 2’FL growth profile, *B. infantis* LH23 strain utilised LNnT, but associated by-products did not support growth of either *B. breve* isolates from infant V2 (Fig 5C). *B. longum* LH206 and LH277, and *B. infantis* LH664 and LH665 grew in the presence of LNnT, however only conditioned media from *B. longum* LH206 generated LNnT-metabolic compounds to support growth of other non-HMO strains in infant V3. We observed moderate growth for *B. pseudocatenulatum* LH656, LH657 and LH663, indicating a cooperative LNnT cross-feeding network within the same microbial ecosystem.

### Metabolite analysis indicates specific metabolic components involved in syntrophic network

We next sought to identify the metabolic compounds involved in these cross-feeding networks for both 2’FL and LnNT HMOs. Using ^1^H-NMR we analysed the metabolic compounds generated by *B. longum* LH206 after growth, using 2’FL as a sole carbon source, and compared that to *B. pseudocatenulatum* LH659 after growth on cell-free supernatant from LH206 (Fig 6A). *B. longum* LH206 produced formate, lactose, galactose, glucose, fucose and acetate in response to growth on 2’FL, and these compounds were reduced after *B. pseudocatenulatum* LH659 growth, suggesting active metabolism (Fig 6A, Fig S5 and Table S9). In addition, LH659 when grown on LH206 conditioned media resulted in production of the metabolite intermediate pyruvate. Cross-feeding experiments using strains from infant V1, indicated that HMO degrader *B. pseudocatenulatum* LH13 and HMO non-degrader *B. longum* LH12 produced similar results (Fig S6).

**Figure 6.**
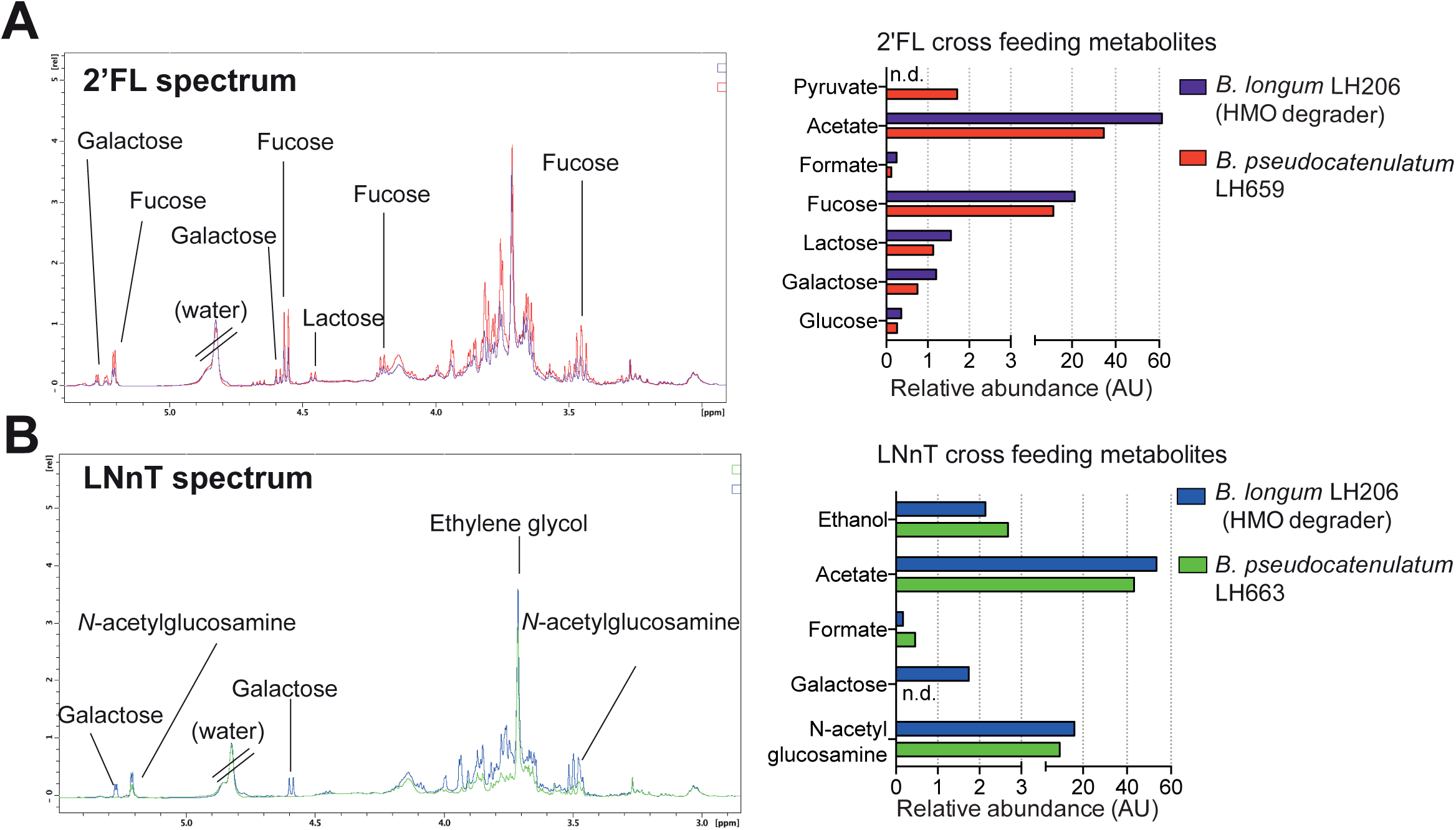
Identification of metabolites involved within cross-feeding. (A)^1^H NMR spectrum to identify metabolites involved in cross-feeding between 2’FL-degrader *B. longum* LH206 and non-degrader *B. pseudocatenulatum* LH659 and semi-quantification of identified metabolites. (E) ^1^H NMR spectrum to identify metabolites involved in cross-feeding between LNnT-degrader *B. longum* LH206 and non-degrader *B. pseudocatenulatum* LH657 and semi-quantification of identified metabolites. n.d indicates metabolites were not detected.

*B. longum* LH206 can also use LNnT for growth, therefore we also profiled cross-feeding metabolites using the non-LNnT degrader, *B. pseudocatenulatum* LH663 from infant V3 (Fig 6B, Fig S7 and Table S9). Similar to the 2’FL cross-feeding experiments, we observed production and consumption of many metabolites; in particular LH206 generated free galactose from LNnT digestion that was no longer detectable in the conditioned media from LH663, indicating galactose had been consumed for growth. There was also an increase in the energy-related compound formate and the end product of fermentation ethanol, suggesting that LH663 was also producing these metabolites in response to growth. In addition, we also examined the metabolic profile of strain *B. longum* LH206 cross-feeding to *B. pseudocatenulatum* LH657 and found additional ethanol production from LH657; but also noted that it consumed acetate, galactose, and N-acetyl glucosamine, by products of LnNT degradation by *B. longum* LH206 (Fig S8). Notably, strain *B. longum* LH206, and not isolate LH277, could digest all HMOs tested, and generated a variety of metabolic by-products that could be consumed by other bifidobacterial strains. Although strain specific, metabolic by-products produced by these HMO degraders, can be used as ‘public goods’ to be shared amongst the *Bifidobacterium* community, thus demonstrating cooperative traits that likely contributes to bifidobacterial survival in the early life infant gut (Fig 7).

**Figure 7:**
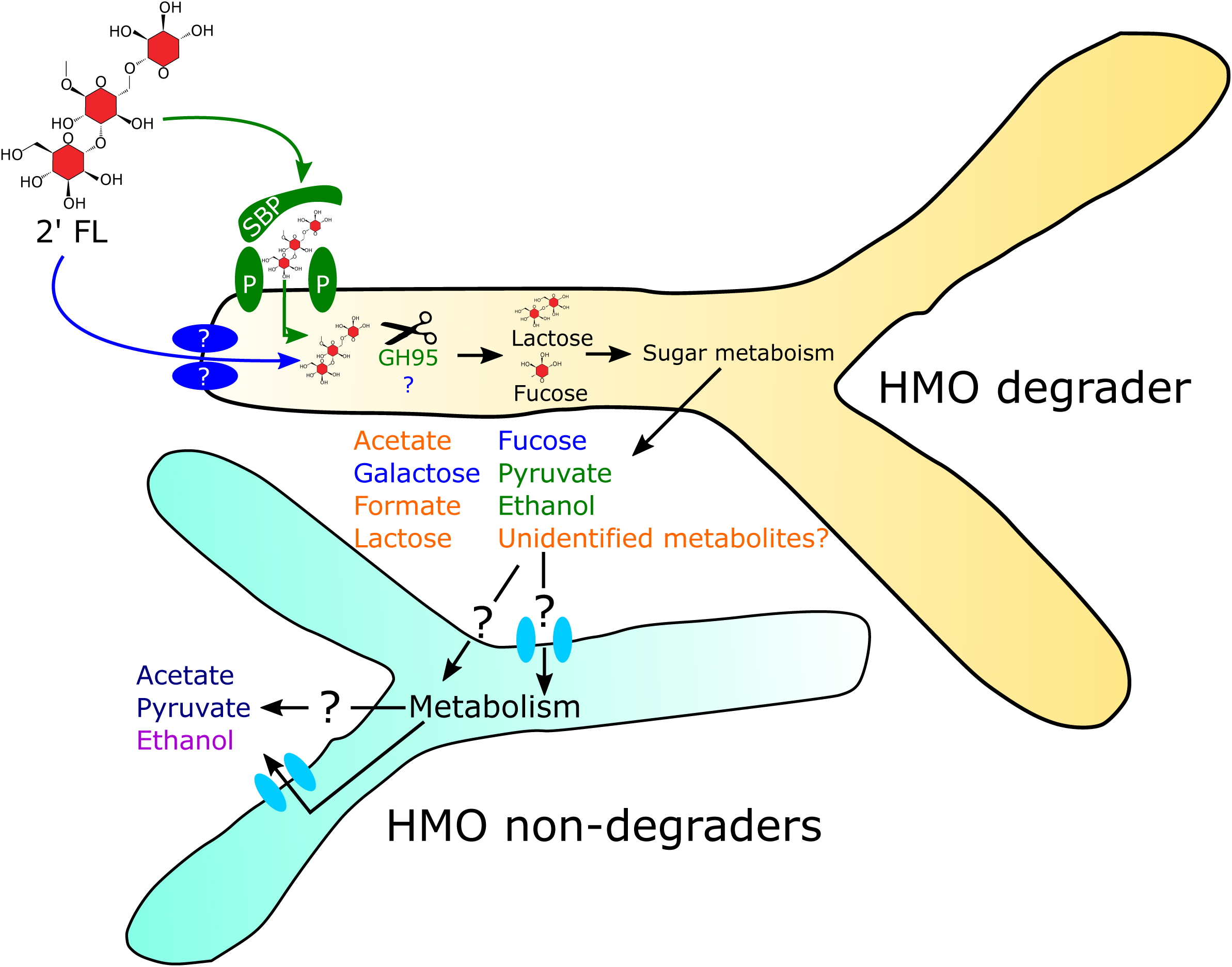
Schematic of the overview for the syntrophic cross-feeding network for HMO 2’FL. We have built a schematic combining the known identified metabolites generated and consumed from 2’FL degradation by *Bifidobacterium* species. Known structures used to import 2’FL into either *Bifidobacterium* isolates LH13 and LH206 (HMO degrader) are denoted in green and blue, respectively. Secreted by-products from 2’FL metabolism are also labelled depending on which strain produced them; LH13 (green), LH206 (blue), or both (orange). Identified substrates generated after cross-feeding to non-HMO utilisers are shown as orange for LH12, or navy for both LH12 and LH659. Additional unidentified mechanisms and metabolites that may be involved in this event are denoted by a question mark (?). Abbreviations: SBP, substrate binding protein; P, permease; GH95, glycoside hydrolase family 95 protein, 2’FL 2’-Fucosyllactose.

## Discussion

*Bifidobacterium* spp. are central players in the early life microbiota and healthy infant development. We show that this genus is present at very high levels in breast fed infants, and that distinct bifidobacterial communities exist within an individual infant ecosystem. Our data indicates a diverse genomic landscape for these individual strains, which links to their ability to thrive on breast milk-associated dietary components i.e. HMOs within a ‘community’ setting. These data highlight the important role that communities of bifidobacteria play in an early life environment, and suggests avenues for development of dietary and microbial-based supplements and therapies to promote infant health.

From three healthy breast-fed infants we isolated a significant number of diverse bifidobacterial strains and species including members commonly associated with the infant microbiota i.e. *B. infantis, B. longum, B. breve* and *B. pseudocatenulatum* (29,30). One proposed method that *Bifidobacterium* are acquired by infants is through vertical transmission from their mother’s vaginal canal during birth as vaginally born infants tend to have a higher proportion of *Bifidobacterium* than those delivered by C-section (31,32). However, horizontal transmission may also occur as the microbiota profile of C-section and vaginally born infants converge after as little as six weeks (33). Both vertical and horizontal transmission of *Bifidobacterium* to the infant gut require the ability to withstand oxygen exposure and passage through the environmental stressors of GI tract i.e. acid and bile exposure (34). In this study, 18 of 19 strains were showed resistance to all the above environmental stressors and this indicates these strains have adapted to allow for effective transmission to the host.

There are distinct genotypic and phenotypic differences between the strains within a single infant bifidobacterial community; however, these differences enable a flexible and cooperative relationship in a breast milk diet (i.e. HMOs) environment that may support dominance of this genus in the wider early life microbiota. HMOs represent a key nutritional component of breast milk, but these complex carbohydrates cannot be directly metabolised by the infant (35). In infants specialised members of the resident gut microbiota allow the breakdown of these compounds, in particular *Bifidobacterium* utilises HMOs and likely contributes to its function as a foundation genus in early life (17,27). Multiple studies have identified genomic clusters for the degradation of these oligosaccharides, including specific clusters for utilisation of specific HMOs (18,21,22). We determined that the genomic arrangement of these clusters exhibits interspecies variability and, consistent with other studies, the presence of these clusters does not always result in a growth phenotype on specified HMOs. For instance, both *B. breve* strains in this study possessed a key GH for fucosylated HMO degradation, but did not grow on 2’FL which could potentially be due to the lack of a second fucosidase (GH29) or appropriate transport genes (20). Furthermore, we identified growth on HMOs in strains lacking known clusters, suggesting a wider variety of novel gene clusters devoted to HMO degradation that could be explored further to provide more mechanistic rationale for development of early life microbiota therapies

Cross-feeding of HMO metabolites between *Bifidobacterium* is a proposed mechanism to allow for non-HMO degraders to survive in the HMO rich environment of the breast-fed infant gut (12,25,36). Recently it has been shown that extracellular sialidases on *B. longum* produce sialylated carbohydrates and free sialic acid to promote *B. breve* growth (24,25); however, there is little evidence suggesting co-operation between non-extracellular HMO degraders. We have identified that by-products of HMO degradation can be utilised by non-degraders from the same infant suggesting sharing of resources within a single host. For example, by products of 2’FL degradation from *B. pseudocatenulatum* strains can support the growth of *B. longum* (Fig 7). Cross-feeding has not been described for *B. pseudocatenulatum* to *B. longum* previously, however, 2’FL has been shown to allow cross-feeding between the intracellular 2’FL degrader *B. infantis* and another microbiota member, *Eubacterium halli*, driven by the 2’FL metabolic end product of 1,2, propanediol (37). Whilst we did not identify this specific product in our system, we did see the consumption of other end products of carbohydrate metabolism including fucose, acetate and pyruvate, which may be sustaining growth in these cross-feeding events. This suggests a more generic sharing of resources between *Bifidobacterium* strains that may also encourage growth of the wider infant gut microbiota including minor, but important members such as *Escherichia coli* (38) and *Staphylococcus* (39), with a mean proportion of 2.5±0.01% and 0.08±0.05% respectively, within our infant microbiotas.

The impact of community sharing of metabolites suggests cooperation between *Bifidobacterium* strains in the infant gut and eludes to the lack of resource competition that has previously been described for identical species within a single ecosystem. It is plausible that due to the abundance of *Bifidobacterium* in early life these cross-feeding events occur specifically within the *Bifidobacterium* community. This advantage is likely key for bifidobacteria survival and ultimate exclusion of potential invading bacteria into the early life microbiota, and suggests *Bifidobacterium* are foundation species that may act as ecosystem engineers, constructing and shaping the gut microbiota in early life. Our results demonstrate that HMO utilisation is strain-specific, and *Bifidobacterium* ultimately have different and complementary carbohydrate degradation mechanisms. Since we examined strains from within a single ecological niche, it is likely that there is shared used of available resources from the infant diet through metabolic cooperation and enables engraftment of *Bifidobacterium* in the infant microbiota. Similar to our findings, recent work has also proposed cooperation amongst multiple *Bifidobacterium* isolates from an SHIME model colon system based on their ability to utilise and share the metabolic components of inulin-type fructans and arabinoxylan oligosaccharides present in the adult diet (40). Despite the concept that similar strains of a specific species tend to compete against each other for shared resources and stable engraftment of new bacteria is enhanced in the absence of other members of the same species (41), our data provides support that positive interactions, like cooperation phenotypes, predominate in single microbial communities and environments (42). In this work, for the first time we have identified altruistic behaviour in *B. longum* strain LH206 to produce a variety of metabolites from HMO degradation that directly promotes the growth of other *Bifidobacterium* strains within the same environment, clearly demonstrating a form of social cooperation between microbes at an individual level. However, what remains to be determined is whether or not there is also mutualistic cooperation occurring, i.e. that LH206 receives a benefit from other strains within the community. Collectively, this research provides new insight into the social behaviour of these proposed foundation species and how they function collectively as a group to maximise nutrient utilisation from breast milk to dominate the infant gut. Determining these interactions with respect to infant diet, will be critical for development of optimal multiple strain/species microbiota therapies to promote early life health.

## Methods

For detailed information see supplementary Materials and Methods.

### Bacterial isolation and strains

Sample collection was in accordance with protocols laid out by the National Research Ethics Service (NRES) approved UEA/QIB Biorepository (Licence no: 11208) and Quadram Institute Bioscience Ethics Committee. Infant faces were isolated on RCM (Oxoid, Hampshire, UK) supplemented with mupirocin and L-cysteine (0.05 mg/mL each, Sigma-Aldrich, Dorset, UK). All *Bifidobacterium* and *Lactobacillus* strains were grown at 37°C in either RCM, de Man Rogosa and Sharpe (MRS) media, or modified MRS (mMRS) with specified carbohydrates in an anaerobic chamber (Don Whitley Scientific, Bingley, UK), unless otherwise specified.

### 16S rRNA gene library preparation and bioinformatics analysis

DNA extraction was performed using the FastDNA Spin Kit for Soil (MPBIO, California, USA) and the V1-V2 region of the 16S rRNA gene was amplified by PCR and run on the Illumina MiSeq platform as previously described (43). Raw reads were initially processed through quality control using FASTX-Toolkit53 maintaining a minimum quality threshold of 33 for at least 50% of the bases. Passed read were then aligned against the SILVA database (44) using BLASTN55 (45) separately for both pairs. All output files were annotated using the paired-end protocol in MEGAN (46).

### Genomic DNA extraction

Bacterial pellets were lysed with lysozyme, Proteinase K, RNase A (all from Roche Molecular Systems, West Sussex, UK), EDTA and Sarkosyl NL30 (Sigma-Aldrich). Samples were purified with three rounds of Phenol:Chloroform:Isoamyl Alcohol (25:24:1) (Sigma-Aldrich) extraction followed by two rounds of extractions with Chloroform:Isoamyl Alcohol (24:1) (Sigma-Aldrich). Genomic DNA pellets were resuspended in 10□mM Tris (pH 8.0) and quantified using Qubit dsDNA BR assay kit (Invitrogen).

### Whole genome sequencing

Isolated DNA was subject to multiplex Sanger Illumina library preparation protocol followed by sequencing using Illumina HiSeq 2500 platform with read length 2 × 125□bp (paired-end reads) and an average sequencing coverage of 60×. Draft genome assemblies were generated using previously described assembly and annotation pipeline (47). Additionally, previously assembled publicly available sequences (n=64) were retrieved online from NCBI Genomes database (48). All genomes were annotated using Prokka v1.10 (49).

### Phylogenetic analysis of whole genomes

General feature format files (GFF) of 83 *Bifidobacterium* strains were used as input for Roary pangenome pipeline v. 3.8.0 to obtain core-genome data (50). The phylogeny was reconstructed from the core-genome alignment generated using MAFFT v 7.305b (51) and subject to cleaning from poorly aligned positions using Gblocks (52,53) and manual curation. Maximum likelihood analysis was performed in Seaview v. 4.0 (54) using PhyML v. 3.1 with 100 bootstrap iterations (55). Python 3 module pyANI with default BLASTN+ settings was employed to calculate the average nucleotide identity (ANI) between the 83 *Bifidobacterium* genomes (56). A cut-off of 95% identity was used for species delineation.

### Functional annotation of genomes

For each genome, all ORFs were submitted to eggNOG-mapper for annotation and classification (57,58). Prediction of HMO clusters was performed by comparing known protein sequences to the draft genomes in this study using local BLAST (45) (e-value<1e^−50^, percentage identify > 70%). HMO clusters were annotated ‘present’ if over 90% of genes were homologous in the cluster. For prediction of GH, ORFs were submitted to the dbCAN web server (59) and the number of GH were calculated. Prophage presence was predicted using PHASTER (60,61).

### Bile salt survival and hydrolysis

To determine *Bifidobacterium* survival in bile, isolates were grown in MRS ± 0.3% unfractionated bovine bile salt (Sigma-Aldrich) as described by (62). Bile salt hydrolyase activity was assessed on MRS agar supplemented with L-cysteine and 0.5% w/v of either taurocholic acid, taurodeoxycholic acid, and sodium glycodeoxycholate bile salt (Sigma-Aldrich). Bile salt precipitation was assessed after a maximum of 96 hours of anaerobic incubation at 37°C.

### Aerotolerance

Each strain was grown aerobically or anaerobically at 37°C and absorbance at 600nm (OD_600nm_) was measured after 48 hours

### HMO utilisation and cross-feeding

*Bifidobacterium* growth in mMRS + 2% (w/v) LNnT or 2’FL (Glycom, Hørsholm, Denmark) was determined using a microplate spectrophotometer. Cell free supernatants (CFS) were prepared by sterile filtration of cultures grown in mMRS + 5% (w/v) LNnT or 2’FL. For cross-feeding, growth in CFS:mMRS (1:1) was monitored every 15 min for 48 in a microplate spectrophotometer (Tecan Infinite F50).

### ^1^H-Nuclear Magnetic Resonance (NMR) Spectroscopy analysis

For functional assessment of *Bifidobacterium* strains, media in which the bacterial cells had been grown, were analysed using ^1^H-NMR Spectroscopy. Media samples were mixed (2:1) with 0.2M sodium phosphate buffer solution (pH 7.4) made in 100% deuterium oxide and 0.01% of sodium 3-(trimethylsilyl) [2,2,3,3,-2H4] propionate 3mM NaN_3_. The mixture was vortexed and centrifuged and transferred to a 5mm outer diameter NMR tube (Wilmad). One-dimensional spectroscopic data were acquired using a 500 MHz NMR spectrometer (Bruker Biospin, Germany) operating at 300 K. A standard one-dimensional NMR pulse sequence with water pre-saturation was applied to acquire spectroscopic data, using 4 dummy scans followed by 64 scans and collected into 24 K data points. ^1^H NMR spectra were manually corrected for phase and baseline distortions and referenced to the TSP signal at δ 0.0, using the TopSpin 3.5 software package (Bruker Biospin, Germany). Spectra from the different bacterial strains grown under different conditions were overlaid in TopSpin and compared for differences. The integrate function was utilised to integrate peaks of interest. Spectral compound libraries (e.g. Human Metabolome DataBase (HMDB), Biological Magnetic Resonance Data Bank) published literature and in-house spectral reference libraries were used to confirm metabolite assignments.

## Supporting information

Supplementary Figures/Tables Summary

## Data availability

All 16S and whole genome data is available at the European Nucleotide Archive, study accession ID PRJEB28188.

## Acknowledgements

We would like to thank the mothers and infants that donated their infant stool samples for this project, and in particular Glycom A/S for the kind donation of purified HMOs: 2’FL and LNnT. The authors would also like to thank Shabhonam Caim and Cristina Alcon-Giner for their technical support with the 16S rRNA pipeline, and Jennifer Ketskemety for help with strain isolations and sequencing prep. This work was funded by a Wellcome Trust Investigator Award (100/974/C/13/Z), and the Biotechnology and Biological Sciences Research Council (BBSRC); Institute Strategic Programme Gut Microbes and Health BB/R012490/1, Institute Strategic Programme Gut Health and Food Safety BB/J004529/1 to LJH, and support of the BBSRC Norwich Research Park Bioscience Doctoral Training Grant (BB/M011216/1, supervisor LJH, student MK). MAEL was funded by a Marie Sklodowska-Curie Individual Fellowship (Project 661594).

## Conflict of interest

All authors declare no conflicts of interest

