## Supplementary Figure 1 for "Infant diet promotes *Bifidobacterium* community cooperation within a single ecosystem"

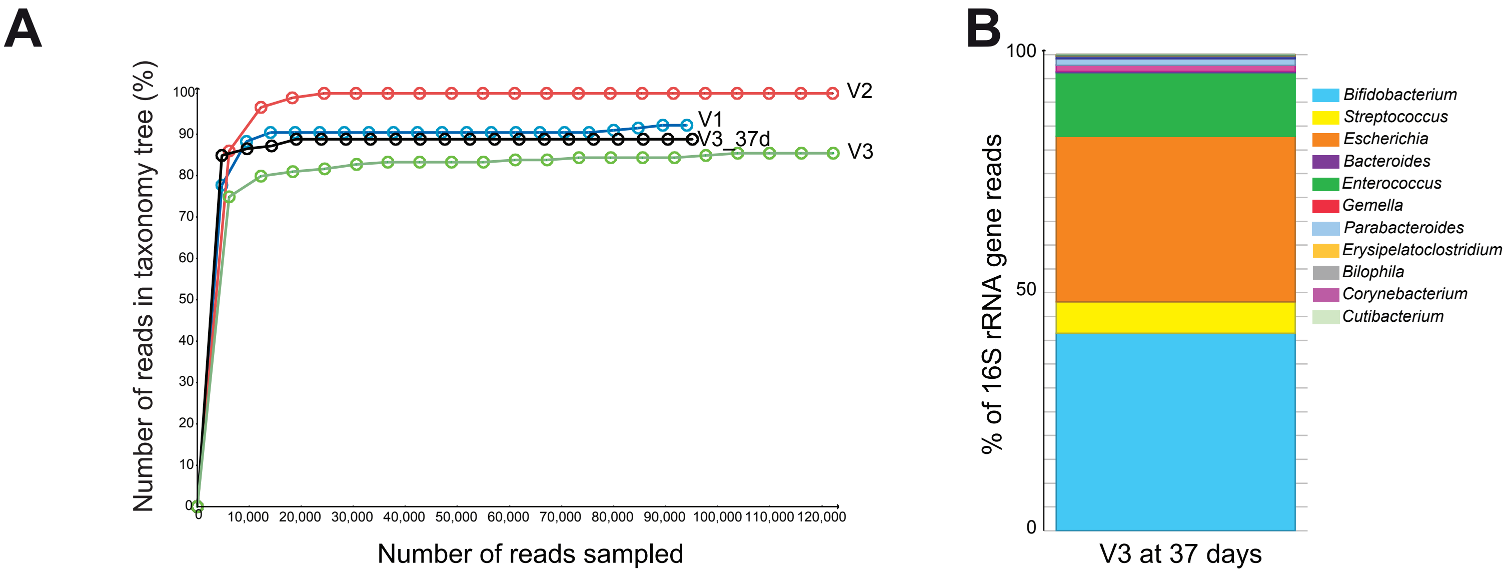


**Supplementary figure 1.** (A) taxonomy refraction curves from 16S rDNA data from infant fecal sample from infant V1, V2, V3 at both 37 and 159 days. (B) 16S rDNA profile of  top 11 Genus identified in infant V3 at 37 days faecal sample shown as a percentage of reads and analysed by MEGAN.
