## Supplementary Figure 2 for "Infant diet promotes *Bifidobacterium* community cooperation within a single ecosystem"

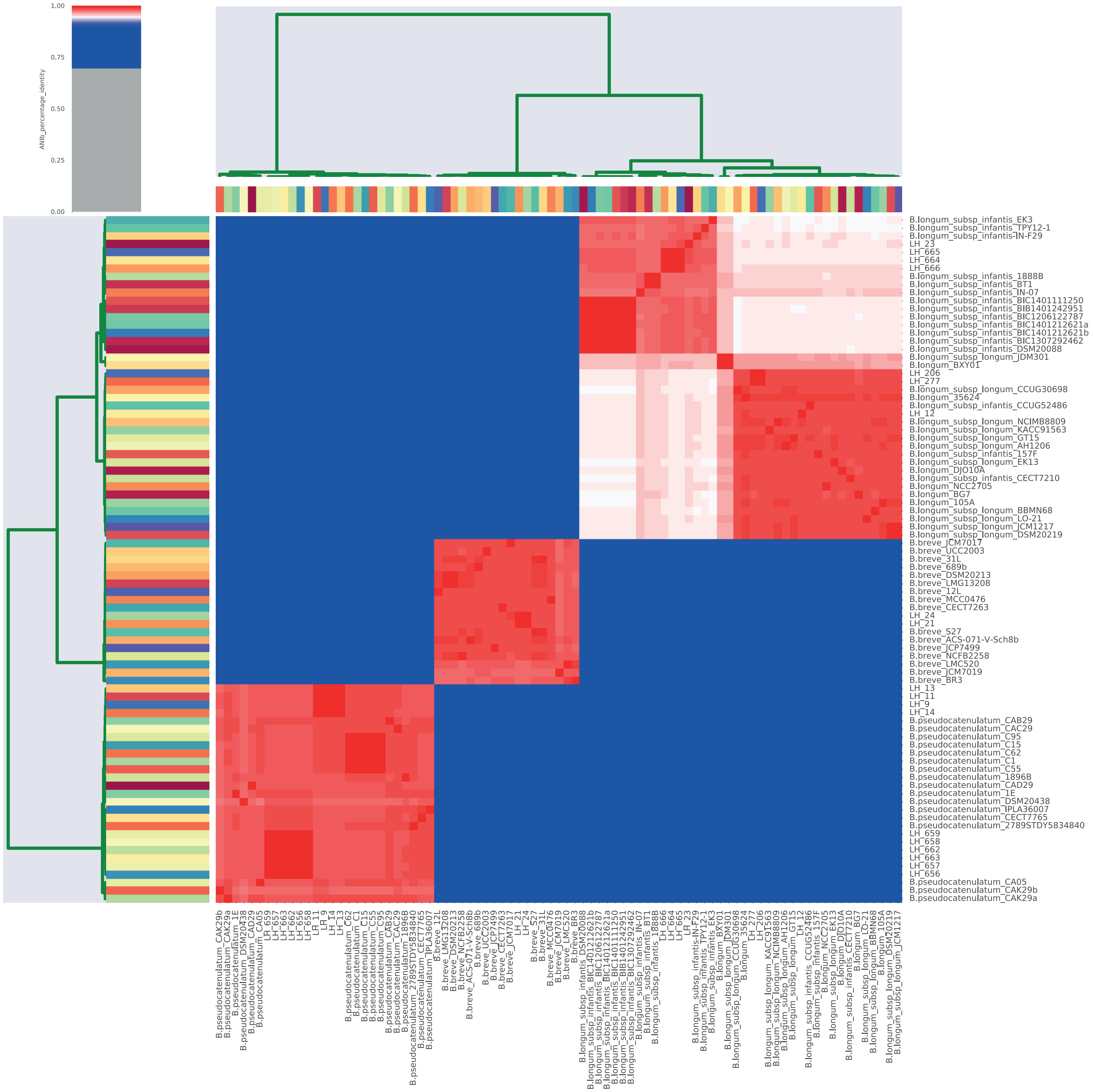


**Supplementary figure 2:** Average Nucleotide Identity (ANI) between 83 *Bifidobacterium* strains. Genetic relatedness above 95% identity (white and red) is used as a cut-off for species delineation.
