## Supplementary Figure 3 for "Infant diet promotes *Bifidobacterium* community cooperation within a single ecosystem"

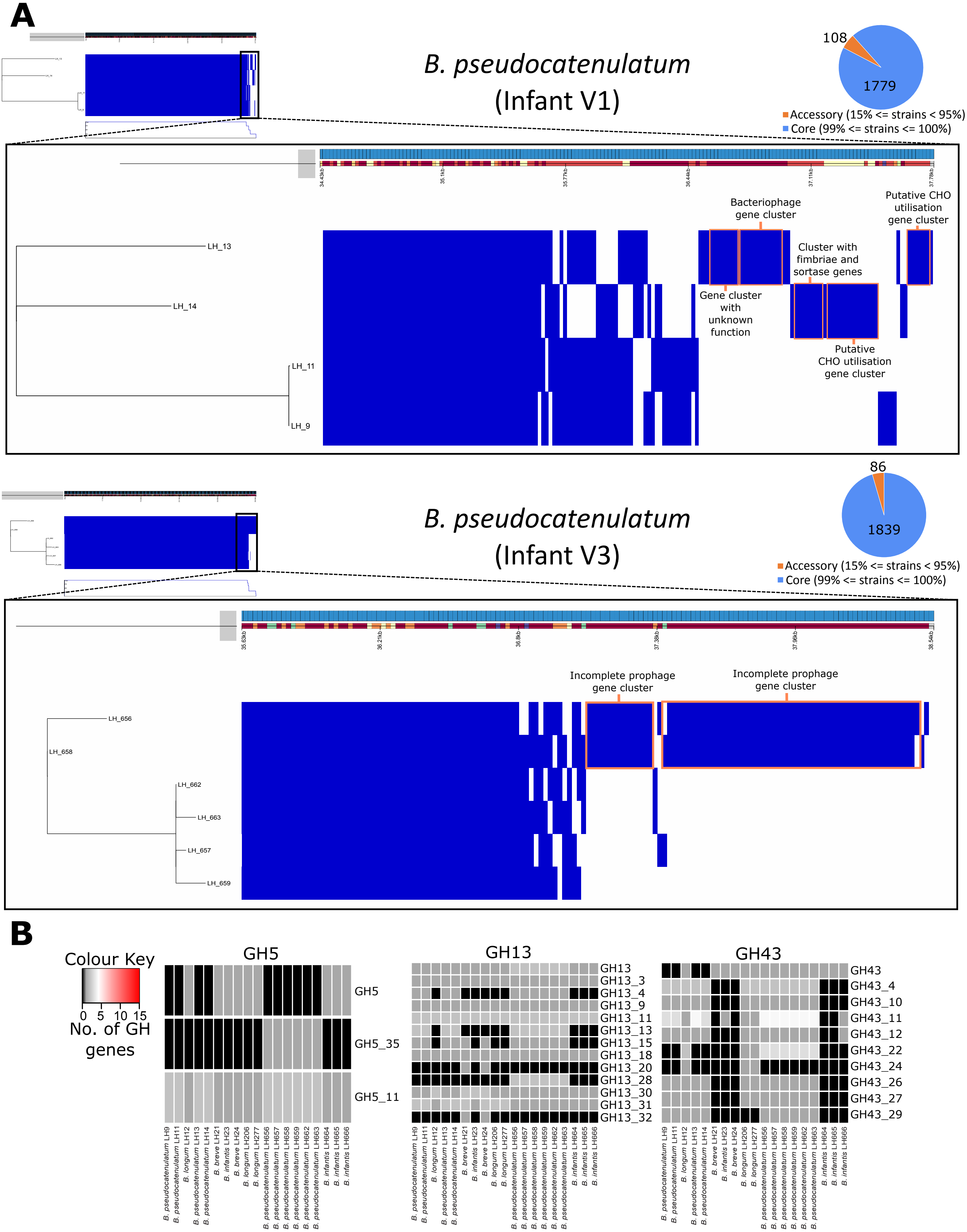


**Supplementary Figure 3:** (A) Visualisation of core and accessory genomes analysis by ROARY for *B. pseudocatenulatum* strains from infant V1 (top panel) and infant V3 (bottom panel). The region of the genome with unique genes is enhanced and gene clusters that are present in only one genome are highlighted in orange. A description of predicted function for each cluster is included. (B) Heatmap indicating the presence of genes classified into glycoside hydrolase subfamilies for all genomes in this study.
