## Supplementary Figure 4 for "Infant diet promotes *Bifidobacterium* community cooperation within a single ecosystem"

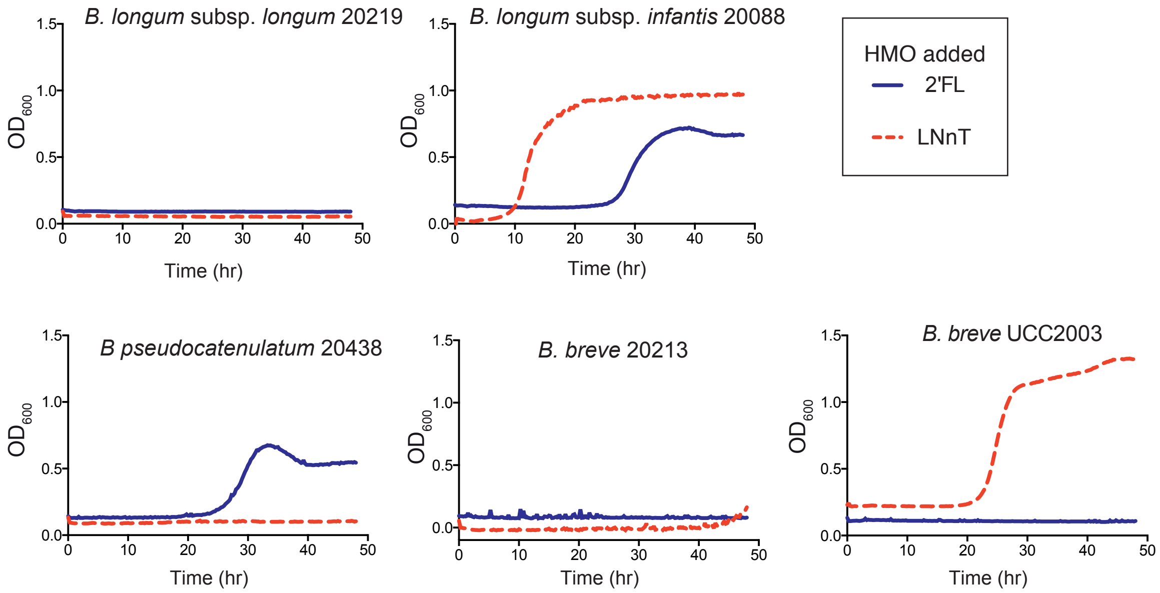


**Supplementary Figure 4:** Growth kinetics of type strains in mMRS with either HMO 2’FL (blue) or LNnT (red) as a sole carbon source. Data shown is a representative graph of three independent experimental repeats, containing the mean from triplicate well measurements.
