## Supplementary Figure 5 for "Infant diet promotes *Bifidobacterium* community cooperation within a single ecosystem"

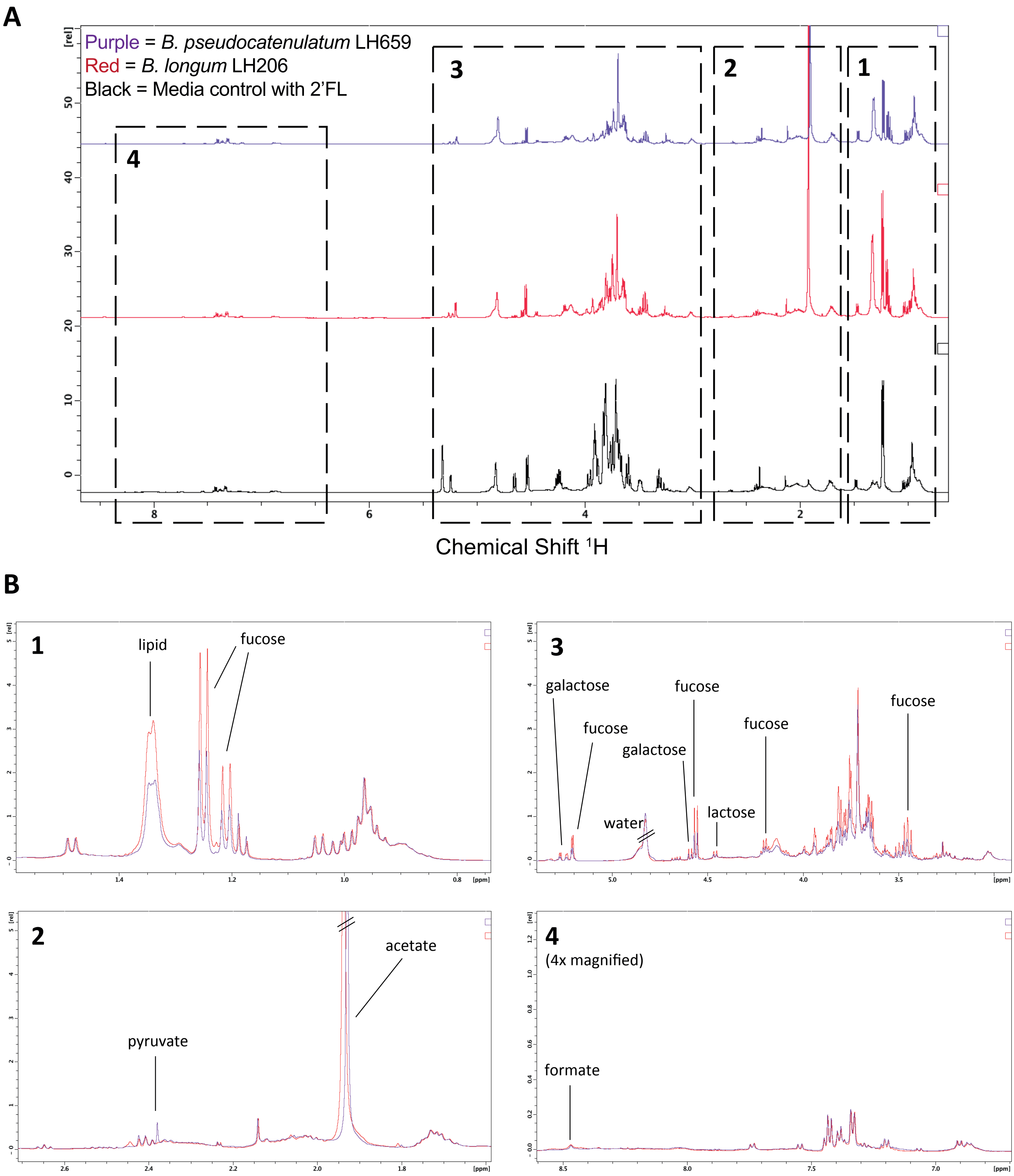


**Supplementary figure 5:** (A)Whole 1H NMR spectrum for 2'FL cross-feeding experiment between Red = *B. longum* LH206 (HMO degrader), Purple = *B. pseudocatenulatum* LH659, and media control supplemented with 2% w/v 2'FL (black) (B) Enlarged overlapping spectrums identified in (A) of metabolites identified after *B. longum* LH206 growth, and the subsequent growth of *B. pseudocatenulatum* LH659 on the conditioned media with annotated metabolites.
