## Supplementary Figure 6 for "Infant diet promotes *Bifidobacterium* community cooperation within a single ecosystem"

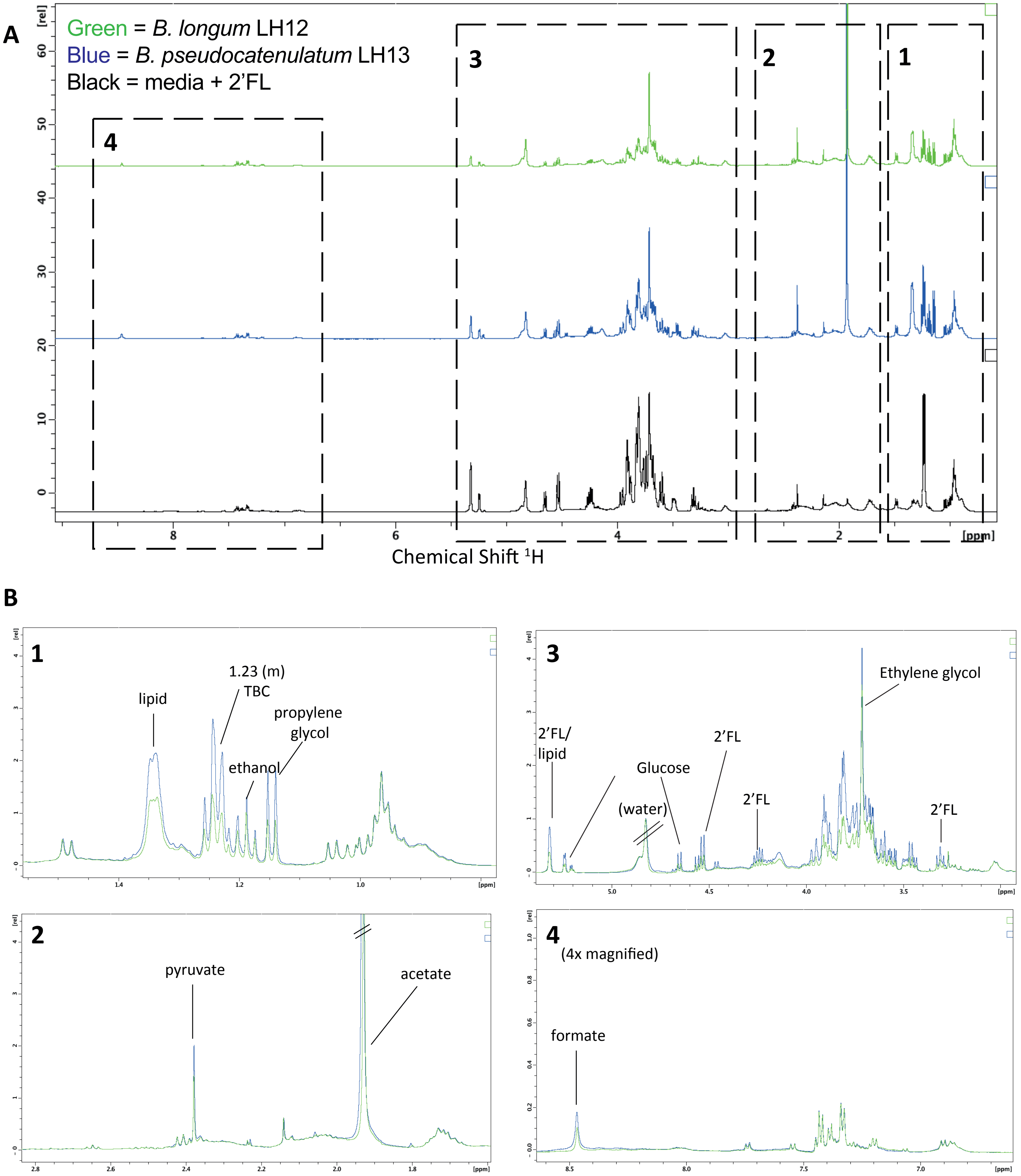


**Supplementary figure 6:** (A) Whole 1H NMR spectrum for LNnT cross-feeding experiment between Blue = *B. pseudocatenulatum* LH13 (HMO degrader), Green = *B. longum* LH13, and media control supplemented with 2% w/v 2'FL (black) (B) Enlarged overlapping spectrums identified in (A) of metabolites identified after *B. pseudocatenulatum* LH12 growth, and the subsequent growth of *B. longum* LH13 on the conditioned media with annotated metabolites.
