## Supplementary Figure 8 for "Infant diet promotes *Bifidobacterium* community cooperation within a single ecosystem"

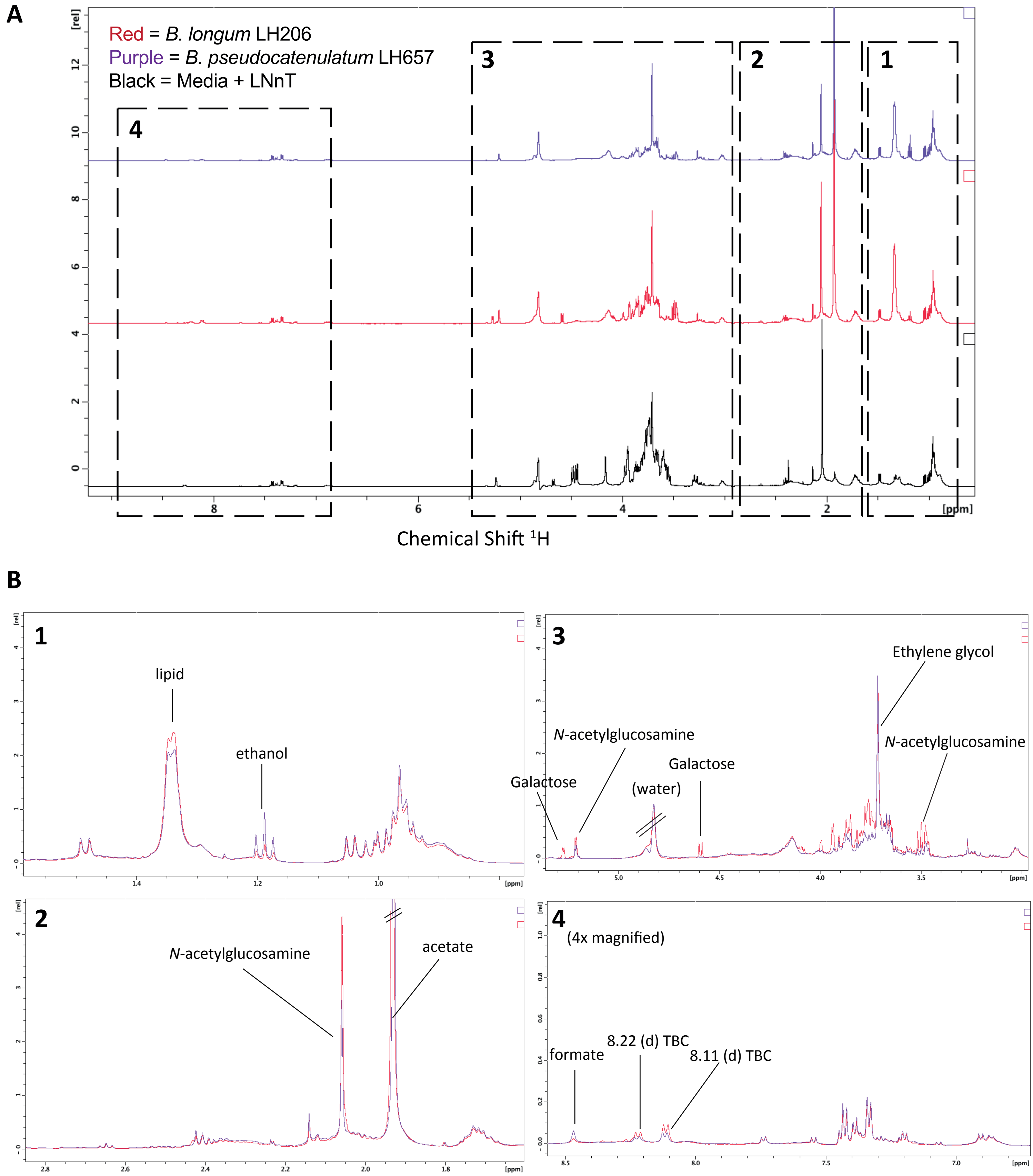


**Supplementary figure 8:** (A) Whole 1H NMR spectrum for LNnT cross-feeding experiment between Red = *B. longum* LH206 (HMO degrader) Purple = *B. pseudocatenulatum* LH657, and media control supplemented with 2% w/v LNnT (black) (B) Enlarged overlapping spectrums identified in (A) of metabolites identified after *B. longum* LH206 growth, and the subsequent growth of *B. pseudocatenulatum* LH657 on the conditioned media with annotated metabolites.
