## Supplementary Methods for "Infant diet promotes *Bifidobacterium* community cooperation within a single ecosystem"

**Bacterial isolation and strains.** Infant faces were collected from freshly soiled nappies and plated on RCM (Oxoid, Hampshire, UK) supplemented with 0.05mg/mL mupirocin and 0.05 mg/mL of L-cysteine (both from Sigma, Dorset, UK) within four hours of collection. All bifidobacteria isolates were grown at 37°C in RCM, de Man Rogosa and Sharpe (MRS) media or modified MRS (mMRS) with specified carbohydrates (including lactose and HMOs) as the main carbon source in an anaerobic chamber (miniMac, Don Whitley Scientific, Bingley, UK), unless otherwise specified. Control and type strains *B. breve* UCC2003, *B. breve* UCC2003DEPS, *B. pseudocatenulatum* DSM20438, *B. pseudocatenulatum* DSM20439, *B. longum* DSM20088, *B. breve* DSM20213, *B. infantis* DSM20219, *Lactobacillus casei* NCIMB4114, and *Escherichia coli* K-12 were also cultured as described above. All samples were collected according Quadram Institute Bioscience Ethics Committee, and sample collection was in accordance with protocols laid out by the National Research Ethics Service (NRES) approved UEA/QIB Biorepository (Licence no: 11208).

**16S rRNA gene library preparation and bioinformatics analysis.** DNA extraction was performed using the FastDNA Spin Kit for Soil (MPBIO, California, USA) following the manufacturer’s instructions, with the exception of extending the bead-beating step to 3 min. Fecal DNA concentration was measured using a Qubit® 2.0 fluorometer (Invitrogen, UK) and normalised to 5ng/mL prior to 16S rRNA gene library preparation. The V1-V2 region of the 16S rRNA gene was amplified using the following primers 5' AGMGTTYGATYMTGGCTCAG and 5' GCTGCCTCCCGTAGGAGT. PCR library prep conditions included an initial 2 min denaturing step at 98°C, followed by 20 cycles of 98°C for 30s, 50°C for 30s, 72°C for 90s, and a final amplification step of 5 min at 72°C run in triplicate on a 96-well T-100 Thermocycler (Bio-Rad, Hertfordshire, UK). PCR products were pooled and purified using AMPure XP beads (Agencourt, Indianapolis, USA) and final quantification was assessed by PicoGreen prior running on the Illumina MiSeq platform with 300 bp paired end reads.

Raw reads were initially processed through quality control using FASTX-Toolkit53 (<http://hannonlab.cshl.edu/fastx_toolkit/index.html>), maintaining a minimum quality threshold of 33 for at least 50% of the bases. Passed read were then aligned against the SILVA database (41) (version: SILVA_132_SSURef_tax_silva) using BLASTN55 (42) (ncbi-blast-2.2.25+; Max e-value 10e-3) separately for both pairs. After performing the BLASTN alignment, all output files were annotated using the paired-end protocol in MEGAN bioinformatics software (43). All raw files have been uploaded to the EBA (European Nucleotide Archive, <https://www.ebi.ac.uk/>) study accession ID PRJEB28188.

**Genomic DNA extraction.** Overnight bacterial cultures (10mL) were grown and used for phenol-chloroform DNA extraction. Briefly, bacterial pellets were re-suspended in 2 ml of 25% sucrose in 10 mM Tris and 1 mM EDTA at pH 8.0. Cells were then treated using 50 μl of 100 mg/ml lysozyme (Roche Molecular Systems, West Sussex, UK). Furthermore, 100 μl of 20 mg/ml Proteinase K (Roche Molecular Systems), 30 μl of 10 mg/ml RNase A (Roche Molecular Systems), 400 μl of 0.5 M EDTA (pH 8.0) and 250 μl of 10% Sarkosyl NL30 (Sigma-Aldrich) were added to the lysed bacterial suspension. The samples were then incubated on ice for 2 hours, and then placed in a 50 °C water bath overnight. The next day, we preformed three rounds of Phenol:Chloroform:Isoamyl Alcohol (25:24:1) (Sigma-Aldrich) extraction using Qiagen MaXtract High Density tubes (Qiagen, Manchester, UK). Followed by two rounds of extractions with Chloroform:Isoamyl Alcohol (24:1) (Sigma-Aldrich) to remove residual phenol before performing ethanol precipitation and wash (in 70% ethanol). Once dry, genomic DNA pellets were resuspended in 300 μl of 10 mM Tris (pH 8.0) and quantified using Qubit dsDNA BR assay kit.

**Whole genome sequencing.** Isolated DNA was subject to multiplex Sanger Illumina library preparation protocol followed by sequencing using Illumina HiSeq 2500 platform with read length 2 × 125 bp (paired-end reads) and an average sequencing coverage of 60×. Draft genome assemblies were generated using previously described assembly and annotation pipeline (44). Additionally, previously assembled publicly available sequences (n=64) were retrieved online from NCBI Genomes database (45). All genomes were annotated using Prokka v1.10 (46) (<https://github.com/tseemann/prokka>). All sequences were uploaded to the ENA under project number PRJEB28188.

**Phylogenetic analysis of whole genomes.** General feature format files (GFF) of 83 *Bifidobacterium*strains were used as input for Roary pangenome pipeline v. 3.8.0 to obtain core-genome data (47). The phylogeny was reconstructed from the core-genome alignment generated using MAFFT v 7.305b (48) and subject to cleaning from poorly aligned positions using Gblocks (49, 50) (http://molevol.cmima.csic.es/castresana/Gblocks_server.html) and manual curation. Maximum likelihood analysis was performed in Seaview v. 4.0 (51) using PhyML v. 3.1 with 100 bootstrap iterations (52). Additionally, genomes of the isolates identified as *B. pseudocatenulatum*in infant individuals were subjected to the analysis with Roary, as described above, to obtain information about their core- and accessory-genome.

Python 3 module pyANI with default BLASTN+ settings was employed to calculate the average nucleotide identity (ANI) between the 83 *Bifidobacterium* genomes (53)(<https://github.com/widdowquinn/pyani>). A cut-off of 95% identity was used for species delineation.

**Functional annotation of genomes.** For each genome, all open reading frames were submitted to eggNOG-mapper (<http://eggnogdb.embl.de/#/app/emapper>) for annotation and classification (54, 55). eggNOG-Mapper annotates each ORF and categorises it based on functional properties. The number of genes in each category was calculated and expressed as a proportion of all OFRs in that genome. A heat map of percentage ORFs in each category was generated using the heatmap.2 package in R.

Prediction of HMO clusters was performed by comparing the protein sequences for each gene in a cluster to the draft genomes described in this study using local BLAST (42) with an e-value cut off of 1e^-50^and percentage identify of 70%. A heat map of the presence or absence of each cluster was generated using the heatmap.2 package in R. A cluster was deemed to be present if more a genome contains homologues of over 90% of genes in the cluster. A cluster was annotated as partially present if genes involved in the enzymatic digestion of HMOs, but not those involved in transport, were present and if no genes were identified then the cluster was annotated as absent.

**Prediction of GH.** For each genomes all ORFs were submitted to the dbCAN web server (<http://csbl.bmb.uga.edu/dbCAN/>, (56)). The dbCAN database uses data from the CAZy database ([http://www.cazy.org](http://www.cazy.org/), (57)) to build HMM models of each GH family and applies these model to predict the presence of GH from submitted protein coding sequences. The number of GH in each GH family was calculated for each genome and a heatmap generated using the heatmap.2 package in R.

**Prediction of prophages.** The draft genomes of LH13 and LH656 were submitted to PHASTER (http://phaster.ca/) to predict the presence of prophage genes in the genome (58, 59). The genomic location of these predicted prophage regions was compared to unique genes identified by ROARY.

**Bile salt survival and hydrolysis**. To determine *Bifidobacterium* survival in bile, isolates were first grown in RCM and then subcultured using a 1:50 dilution into MRS ± 0.3% unfractionated bovine bile salt (Sigma-Aldrich), as described by (60). After 48h of stationary growth in an anaerobic chamber at 37°C OD_600nm_ using the Benchmark Plus microplate spectrophotometer (Bio-Rad) for both conditions. For the MRS plate the mean blank OD_600nm_value was 0.1585 and for the anaerobic plate it was 0.1825. Data shown is mean values from three experimental repeats.To assess bile salt hydrolyase activity, overnight cultures were spotted (3mL) onto MRS plates supplemented with L-cysteine and 0.5% w/v of either taurocholic acid, taurodeoxycholic acid, and sodium glycodeoxycholate bile salt (Sigma-Aldrich). Bile salt precipitation was assessed after a maximum of 96 hours of anaerobic incubation at 37°C. For both assays, uninoculated MRS media was used as a control.

**Aerotolerant assay.** MRS media inoculated with a 1:50 dilution of each strain that had been grown aerobically or anaerobically at 37°C for 48 hours stationary. The average absorbance of un-inoculated MRS media was based on was subtracted from all OD_600nm_ readings, as described in (61). For the aerobic plate the blank OD_600nm_ value was 0.1507 and for the anaerobic plate it was 0.1549. Data shown is mean values from three experimental repeats.

**HMO utilisation and cross-feeding.** Growth kinetics of the 19 novel *Bifidobacterium* isolates and control strains using either LNnT or 2’FL (Glycom, Hørsholm, Denmark) as the soul carbon source (2%w/v) in mMRS was determined using a microplate spectrophotometer. For cross-feeding experiments, HMO-users were grown in mMRS and LNnT or 2’FL in an excess of 5%w/v, to ensure the carbon source was not the limiting factor for overnight growth. Cultures were then centrifuged at 4°C for 10 minutes at 4,000 rpm and the supernatant was sterile filtered (0.22mm filter) twice, prior to being used for growth with strains that had been identified to be unable to degrade the HMOs. Spent media was then added in a 1:1 ratio with mMRS media and then inoculated with the specified strains and monitored for growth by a microplate spectrophotometer (Tecan Infinite F50) with agitation every 15 minutes for 48 hours.

**^1^H-Nuclear Magnetic Resonance (NMR) Spectroscopy analysis.** For functional assessment of *Bifidobacterium* strains, media in which the bacterial cells had been grown, were analysed using ^1^H-NMR Spectroscopy. Media samples were prepared for analysis, by transferring 400μL of cell media into a microtube containing 200μL of 0.2M sodium phosphate buffer solution (pH 7.4) made in 100% deuterium oxide (which was needed for the field lock of the NMR spectrometer). The buffer solution also contained 0.01% of sodium 3-(trimethylsilyl) [2,2,3,3,-2H4] propionate (TSP) as an internal reference standard for calibration of acquired spectral profiles, and 3mM NaN3 as a preservative. The mixture was vortexed and centrifuged for 10 seconds followed by transfer into a 5mm outer diameter NMR tube (Wilmad).

One-dimensional spectroscopic data were acquired using a 500 MHz NMR spectrometer (Bruker Biospin, Germany) operating at 300 K. A standard one-dimensional NMR pulse sequence with water pre-saturation was applied to acquire spectroscopic data, using 4 dummy scans followed by 64 scans and collected into 24 K data points. ^1^H NMR spectra were manually corrected for phase and baseline distortions and referenced to the TSP signal at *δ* 0.0, using the TopSpin 3.5 software package (Bruker Biospin, Germany). Spectra from the different bacterial strains grown under different conditions were overlaid in TopSpin and compared for differences. The integrate function was utilised to integrate peaks of interest. Spectral compound libraries (e.g. Human Metabolome DataBase (HMDB), Biological Magnetic Resonance Data Bank) published literature and in-house spectral reference libraries were used to confirm metabolite assignments.
