## Supplementary Figures/Tables Summary for "Infant diet promotes *Bifidobacterium* community cooperation within a single ecosystem"

STable_1.xlsx - Metadata of infants use, and strains identified in this study
STable_2.xlsx -  No of reads for each genus identifed by 16s rRNA sequencing
STable_3.xlsx -  Details of bile salt hydrolases genes for genomes in this study
STable_4.xlsx -  Results from bile salt hydrolysis assay
STable_5.xlsx -  Data used to generate figure 1A
STable_6a.xlsx -  Details of accessory genes for B. pseudocatenulatum strains from infant V1
STable_6b.xlsx -  Details of accessory genes for B. pseudocatenulatum strains from infant V3
STable_7.xlsx -  Details of partial prophage identifed in stains LH13 and LH656
STable_8.xlsx -  Presence of extracellular sialidase genes for genomes in this study
STable_9.xlsx -  1H NMR metabolite peak annotation for cross-feeding experiments														
Supp_1.pdf -  Figure containing Additional 16s data (taxonomy refraction curves and assignment) 
Suppl_2.pdf -  Figure showing Average nucleotide identity between 83 publised bifidobacterial genomes and genomes isolated in this study
Suppl_3_30072018.pdf -  Figure showing Plots showing detail of accessory genomes of B. pseudocatenulatum strains from infant V1 and V3. 
Suppl_4.pdf -  Figure showing Growth curves of reference strains on HMOs
Suppl_5_12072018.pdf -  Figure showing Whole 1H NMR spectrum for 2’FL cross-feeding experiment for infant V1 strains
Suppl_6_12072018.pdf -  Figure showing Whole 1H NMR spectrum for LNnT cross-feeding experiment for infant V1 strains
Suppl_7_12072018.pdf -  Figure showing Whole 1H NMR spectrum for 2’FL cross-feeding experiment for infant V3 strains
Suppl_8_12072018.pdf -  Figure showing Whole 1H NMR spectrum for LNnT cross-feeding experiment for infant V3 strains
Supplementary Methods.docx - Detailed methods
